# A comprehensive pharmacological survey across heterogeneous patient-derived GBM stem cell models

**DOI:** 10.1101/2024.11.27.625719

**Authors:** Richard J.R. Elliott, Peter W.K. Nagle, Muhammad Furqan, John C. Dawson, Vanessa Smer-Barreto, Diego A. Oyarzún, Aoife McCarthy, Alison F. Munro, Camilla Drake, Gillian M. Morrison, Steven M. Pollard, Michael Marand, Daniel Ebner, Val Brunton, Margaret C. Frame, Neil O. Carragher

**Affiliations:** Cancer Research UK Scotland Centre (Edinburgh), Institute of Genetics and Cancer, University of Edinburgh, Western General Hospital, Edinburgh, UK; School of Informatics, University of Edinburgh, 10 Crichton St, Edinburgh, EH8 9AB, UK; School of Biological Sciences, University of Edinburgh, Max Born Crescent, Edinburgh, EH9 3BF, UK; Institute of Regeneration and Repair, Cancer Research UK Scotland Centre, University of Edinburgh, EH16 4UU, U.K.; Nuffield Dept. Of Medicine, University of Oxford, Old Road Campus, Oxford OX3 7BN, U.K.

## Abstract

Despite substantial drug discovery investments, the lack of any significant therapeutic advancement in the treatment of glioblastoma (GBM) over the past two decades calls for more innovation in the identification of effective treatments. The inter-and intra-patient heterogeneity of GBM presents significant obstacles to effective clinical progression of novel treatments by contributing to tumour plasticity and rapid drug resistance that confounds contemporary target directed drug discovery strategies. Phenotypic drug screening is ideally suited to heterogeneous diseases, where targeting specific oncogenic drivers have been broadly ineffective. Our hypothesis is that a modern phenotypic led approach using disease relevant patient-derived GBM stem cell systems will be the most productive approach to identifying new therapeutic targets, drug classes and future drug combinations that target the heterogeneity of GBM. In this study we incorporate a panel of patient-derived GBM stem cell lines into an automated and unbiased ‘Cell Painting’ assay to quantify multiple GBM stem cell phenotypes. By screening several compound libraries at multiple concentrations across a panel of patient-derived GBM stem cells we provide the first comprehensive survey of distinct pharmacological classes and known druggable targets, including all clinically approved drug classes and oncology drug candidates upon multiple GBM stem cell phenotypes linked to cell proliferation, survival and differentiation. Our data set representing, 3866 compounds, 2.2million images and 64000 datapoints is the largest phenotypic screen carried out to date on a panel of patient-derived GBM stem cell models that we are aware of. We seek to identify agents and target classes which engender potent activity across heterogenous GBM genotypes and phenotypes, in this study we further characterize two validated target classes, histone deacetylase inhibitors and cyclin dependent kinases that exert broad and potent effects on the phenotypic and transcriptomic profiles of GBM stem cells. Here we present all validated hit compounds and their target assignments for the GBM community to explore.

## Introduction

Glioblastoma (GBM), a Grade IV astrocytoma, is a highly complex disease that remains difficult to treat. Mortality rates have increased over the last 30 years, especially in the over 60 age group, as incidence rates outpace survival improvements [1]. GBM is therefore the most common and aggressive primary malignant tumour of the brain and represents an urgent area of unmet medical need. Since the introduction of post-surgical, concomitant radiotherapy and adjuvant temozolomide in 2005 (the ‘Stupp protocol’), which increased median overall survival (mOS) by ∼3 months[2], a lack of effective, novel treatments for GBM have been introduced into broad clinical use and 5-year survival rates from time of diagnosis remain at less than 5%[3]. Clinical and scientific research over the last 20 years has revealed a complex landscape of remarkable intra- and inter-tumoral heterogeneity associated with the highly dynamic nature of GBM molecular biology [4, 5{Verhaak, 2010 #312{Brennan, 2013 #313, 6, 7]. The complex biology driving the disease is further exacerbated by the addition of GBM stem cells (GSC), driving self-renewal and resistance mechanisms[8-13]. The self-renewing capacity of GSCs is an inherent feature of GBM to which the tumour microenvironment, or the GBM stem cell niche, contributes by supporting adaptive signal rewiring, heterogeneous tumour evolution and therapeutic resistance[14, 15]. Phospho-proteomic studies performed on patient derived GBM model systems demonstrate rapid rewiring of pathway signalling at the post-translational pathway level following effective inhibition of some of the most compelling drug targets such as the mTOR pathway, that is hyperactivated in approximately 90% of GBM[16]. GBM plasticity and heterogeneity thus confound modern target-based drug discovery strategies contributing to drug resistance and relapse, hence the targeting of clear oncogenic drivers of GBM, such as EGFR amplification/mutation or RAS, PI3K signalling, has not translated into significant efficacy in GBM clinical management. Emerging single-cell transcriptomic[17], proteomic[18] and image-based phenotypic screening[19] technologies combined with improved patient-derived GSC models[20] provide new opportunities to understand and overcome disease heterogeneity. New hope for the development of novel therapeutic classes and drug combinations that may overcome some if this heterogeneity and support new personalized treatments strategies for GBM is provided from the recent phase 2 ROAR basket clinical trial study evaluating a combination of a BRAF targeting agent (dabrafenib) and the MEK inhibitor (trametinib) which includes a cohort of 45 high grade glioma (HGG, including GBM) patients with BRAF V600E mutation. The primary endpoint of investigator-assessed overall response rate was 33% in the HGG cohort. Secondary endpoints of median duration of response (DoR), progression-free survival (PFS) and mOS in the HGG cohort was; 31.2, 5.5 and 17.6 months respectively[21]. Emerging case reports from both adult and pediatric patients also point to the benefit of treating BRAF mutation-positive glioblastoma and gliomas with the dabrafenib and trametinib combination[22-25]. However, BRAF V600E mutation can be found in approximately only 3% of GBM patients therefore, while such personalized treatments offer hope, the challenge is addressing the majority of patients which do not fall into molecular subtypes defined by a well-established druggable target or pathway, or are refractory to such personalized treatments.

The drug development of imidazotetrazines, such as temozolomide, extended from chemocentric and phenotypic-led collaborative research exploring the *in vitro* and *in vivo* anti-cancer properties of small molecules, dacarbazine and mitozolomide (azolastone)[26] emphasizing the value of incorporating complex tumour biology into early stage drug design and selection. Phenotypic drug discovery (PDD), defined as the identification of hit or lead compounds prior to target identification[27], has historically contributed more than target-based drug discovery (TDD), in terms of first-in-class drugs approved by the FDA since 1999[28, 29], yet TDD has dominated drug discovery approaches in oncology over the same period of time[30]. While TDD and PDD are complimentary drug discovery strategies, PDD can offer some advantages over target-based methods, as it allows discovery of potential therapeutics using complex, biological models of poorly understood heterogeneous disease states[31, 32] and can reveal novel unprecedented therapeutic classes such as the first HDAC inhibitor (vorinostat), molecular glues (lenalidomide), spicing modulators (risdiplam)[28, 33] and novel conformation-selective binding modes-of-action for existing targets (NXP900/eCF506)[34]. This ‘target-agnostic’ approach has been greatly assisted by hardware advances in high-content microscopy, the development of advanced image analysis software tools and improved phenotypic profiling techniques, such as Cell Painting and Artificial Intelligence/Machine Learning (AI/ML) methods that help classify cell phenotypes and elucidate drug mechanism-of-action[35-38]. Recent advances in multiparametric high-content imaging enables the generation of a phenotypic fingerprint for every chemical (or genetic) perturbation to support the new discipline of ‘phenotypic (or morphometric) profiling’ that quantifies functional similarities and dissimilarities between drug mechanism-of-action (MOA) within intact cell-based assay systems[35, 39-41]. The Cell Painting assay multiplexes six fluorescent dyes, imaged in five spectral channels to reveal eight broadly relevant cellular components or organelles and can be combined with the development of bespoke automated image analysis pipelines to segment individual cells and calculate thousands of morphological feature-based measurements per cell (e.g. various measures of size, shape, texture, intensity and many others) to produce a data-rich profile that is suitable for the detection of subtle cellular phenotypes which can be linked to biological outcomes[42]. Here we describe the adaptation and optimization of Cell Painting for profiling compound MOA and target-class across a panel of 6 genetically distinct GSC lines obtained from the Glioma Cellular Genetics Resource (https://github.com/GCGR). Using this approach we provide a comprehensive pharmacological audit of how specific target, pharmacological and chemical classes influence patient-derived GBM cellular phenotypes. We present a large-scale application of this technique using chemically diverse libraries containing FDA approved drugs, or Phase I passed clinical candidates and selective chemical probe compounds against known oncology targets. We identify multiple diverse pharmacological and target classes effective across our genetically distinct panel of patient-derived GSCs and present all validated screening hits and their target assignments to the GBM research community. These results support our hypothesis that an unbiased phenotypic led approach, using disease relevant patient-derived GSC models and exploitation of deep phenotyping and drug mechanism-of-action profiling technologies is a productive approach to identifying new targets, drug classes and future drug combinations that address the heterogeneity of GBM.

## Results

### Glioblastoma Stem Cell Models

Patient-derived GSCs were obtained from the Cancer Research UK Glioma Cellular Genetics Resource, and 6 GSC lines were initially selected to cover common transcriptomic subtypes based on their RNA-Seq subtype classification (classical (CLA), mesenchymal (MES), proneural (PRO), Supplementary Figure 1A). GSC’s were cultured on laminin-1 coated substrates, under defined media conditions that have previously been demonstrated to maintain their stem cell-like properties[43]. Initial studies demonstrate compatibility with high-content imaging (i.e., flat and adherent morphology) and cell number was optimised for 384-well microtiter plate formats to enable high-throughput drug screening (Supplementary Figure 1B-D). Given the generally slow growth rates of the GSCs relative to standard cancer cell line cultures, compound exposure over 72 hours during screening was selected to ensure at least one population doubling in all lines. Short-tandem repeat (STR) profiling was carried out on each GSC line to create reference sequences and all cell lines were expanded and cell stocks banked in liquid nitrogen, to facilitate screening with consistent low passage numbers ensuring minimal biological drift across primary screening and secondary hit validation assays. In order to begin characterising these cell lines, prior to phenotypic screening we examined basal protein expression and post-translational pathway activation status by Reverse Phase Protein Array (RPPA) and cytokine array in 2-dimensional (2D) and 3-dimensional (3D) cell culture conditions. Basal levels of protein and phospho-epitope abundance (in 2D and 3D culture) show functional enrichment of signalling pathways in MAPK, ErbB, VEGF, PI3K-AKT, focal adhesion, EGFR inhibitor resistance and PD-1 checkpoint in cancer (Supplementary Figure 2). The cytokine array profiling further demonstrates enrichment in the secretion of GBM associated chemokine proteins involved in tumour progression, invasion, angiogenesis (IL-6, MMP-7, TIMP1, VEGF), and tumour associated neutrophils (TANs; CCL2, IL-8/CXCL8)[44] including factors previously shown to contribute to GSC subtype trans-differentiation (PRO to MES transition)[45]. (Supplementary Figure 2A-D). Notably the six GSC lines representing three pairs of the common GBM transcriptomic sub-types (CLA, MES and PRO) selected for our study do not cluster into any specific subgroups based on basal protein expression or pathway activation status. Interestingly, high levels of the pro-inflammatory cytokine, interleukin-8 (CXCL8, Supplementary Figure 2C, D) has recently been reported as a druggable target in GBM (*via* humanised anti-IL8 antibody) with an anti-PD-1 combinational blockade[46]. Overall, this pre-screening profiling indicates our proposed GSC drug screening models are representative of typical GBM signalling networks observed in patient samples, express known druggable pathways while capturing the expected broad heterogeneity of GBM tumours at the molecular, protein and morphological levels in both 2D adherent and 3D cell culture.

### GSC Cell Painting Assay Development

Previously, we have successfully adapted the Cell Painting protocol[35, 47] to explore drug MOA across cancer cell line panels from diverse lineages including, breast and oesophageal adenocarcinoma[19, 48, 49]. In the current study, we adapt the protocol to our panel of patient-derived GSCs and incorporate live mitochondrial staining prior to fixation and staining with the remaining Cell Painting reagents (Figure 1A, B). It is our understanding that this study describes the first application of phenotypic profiling using the Cell Painting assay to a panel of patient-derived GSC models. Images from each cell line were then used to build a customized CellProfiler[50] image analysis pipeline and confirm appropriate image segmentation and performance of high content analysis. CellProfiler analysis quantified up to 1006 cellular features from each GSC and principal component analysis was carried out on median aggregated well level data using the StratoMineR HCA platform (stratominer.com). Nuclei Count features were also extracted to assess reproducibility of cell seeding in 384-well format and calculate the percentage cell survival following each compound treatment including signal-to-noise assessment between negative control (0.1% (v/v) dimethylsulfoxide, DMSO) and cytotoxic positive control compounds (paclitaxel, staurosporine). These assay validation studies confirm assay reproducibility and signal-to-noise in 384-well plate formats is suitable for high-throughput screening based on industry-standard recommendations[51] (Supplementary Figures 1C & Supplementary Figure 3).

**Figure 1.**
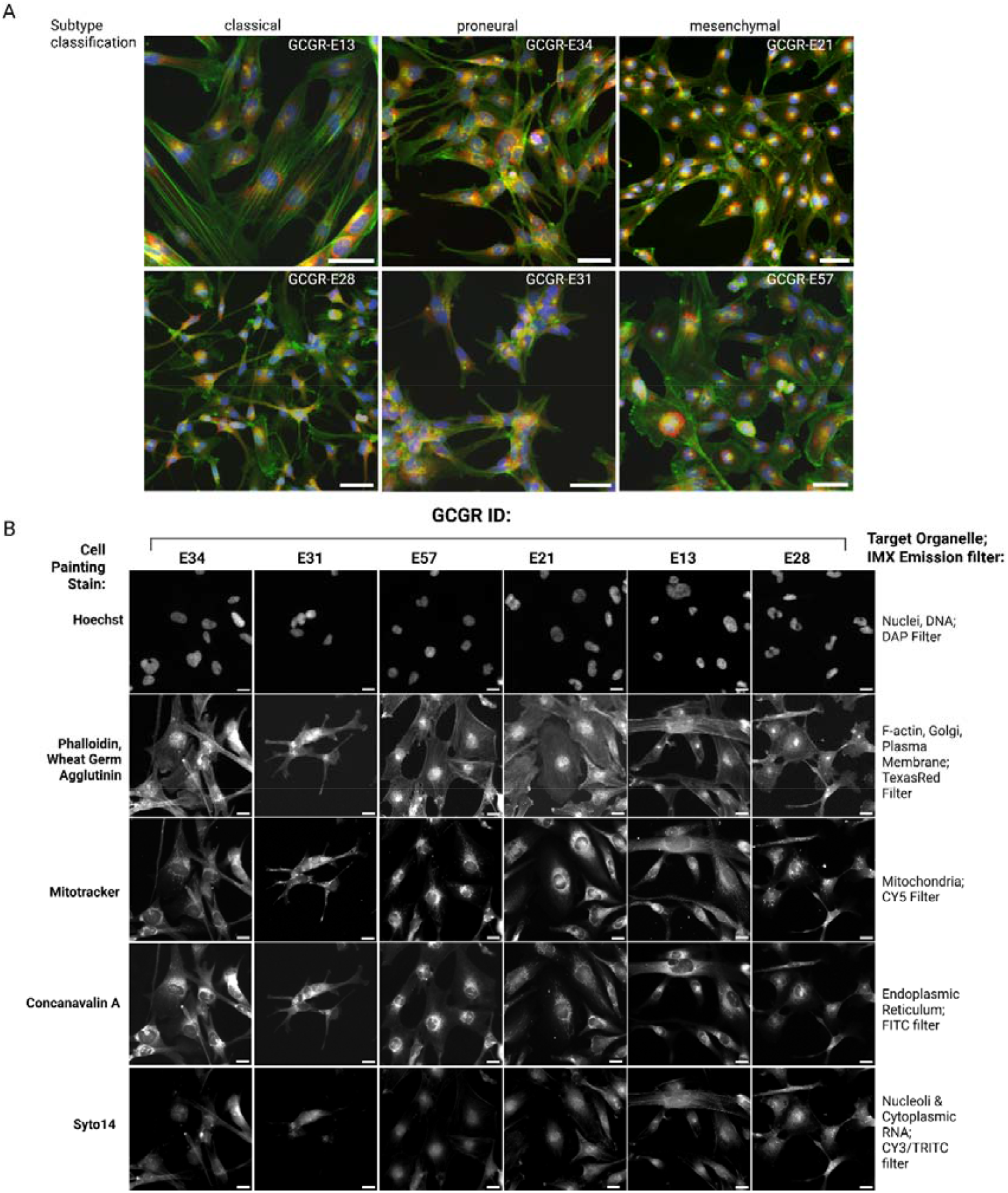
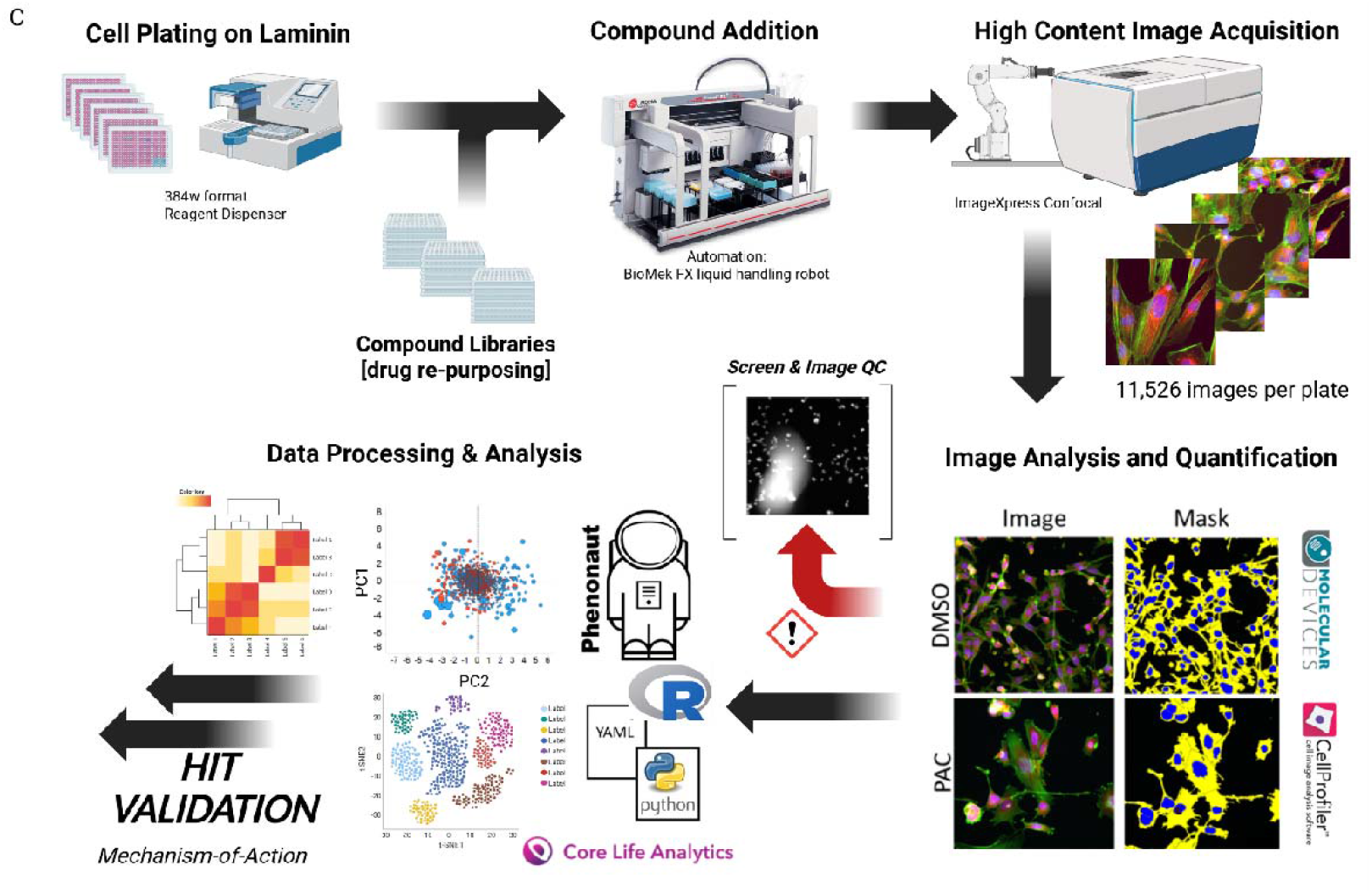
High content phenotypic assay screening on GSC panel. A. Representative images of Cell Painting across GSC panel (Hoechst (blue), Phalloidin/WGA (green), Endoplasmic reticulum (red). Scale bar 50µM. B. Representative images of individual cell painting stains, target organelle and corresponding filter across GSC panel. Scale bar 20µM. C. Established high content screening, image and analysis workflow, followed by subsequent hit validation and exploration.

Our phenotypic screening campaign performed across the panel of six GSC lines incorporated multiple small molecule compound libraries screened against varied concentrations (ranging from 3-10,000nM) (see Table 2 methods section for specific concentrations for each library). Compound libraries included the LOPAC (1280 compounds, Merck), Prestwick FDA (1280 compounds), TargetMol L2110 (330 compounds), Structural Genomics Consortium Kinase Chemogenomic set (KCGS – 295 compounds)[52]) and a custom chemogenomic library set (789 compounds, ‘Comprehensive anti-Cancer small Compound Library, or C3L) (Table 1, Materials & Methods). The C3L library was created with a focus on chemical and known oncology target diversity to functionally evaluate known oncology targets across phenotypic screening assays[53]. The majority of these commercially available compounds (across all library sets) have been through phase I clinical trials. It is our view that this approach presents the potential for discovering novel drug repurposing and drug combination opportunities in addition to identifying new therapeutic targets not previously explored in GBM.

**Table 1.**
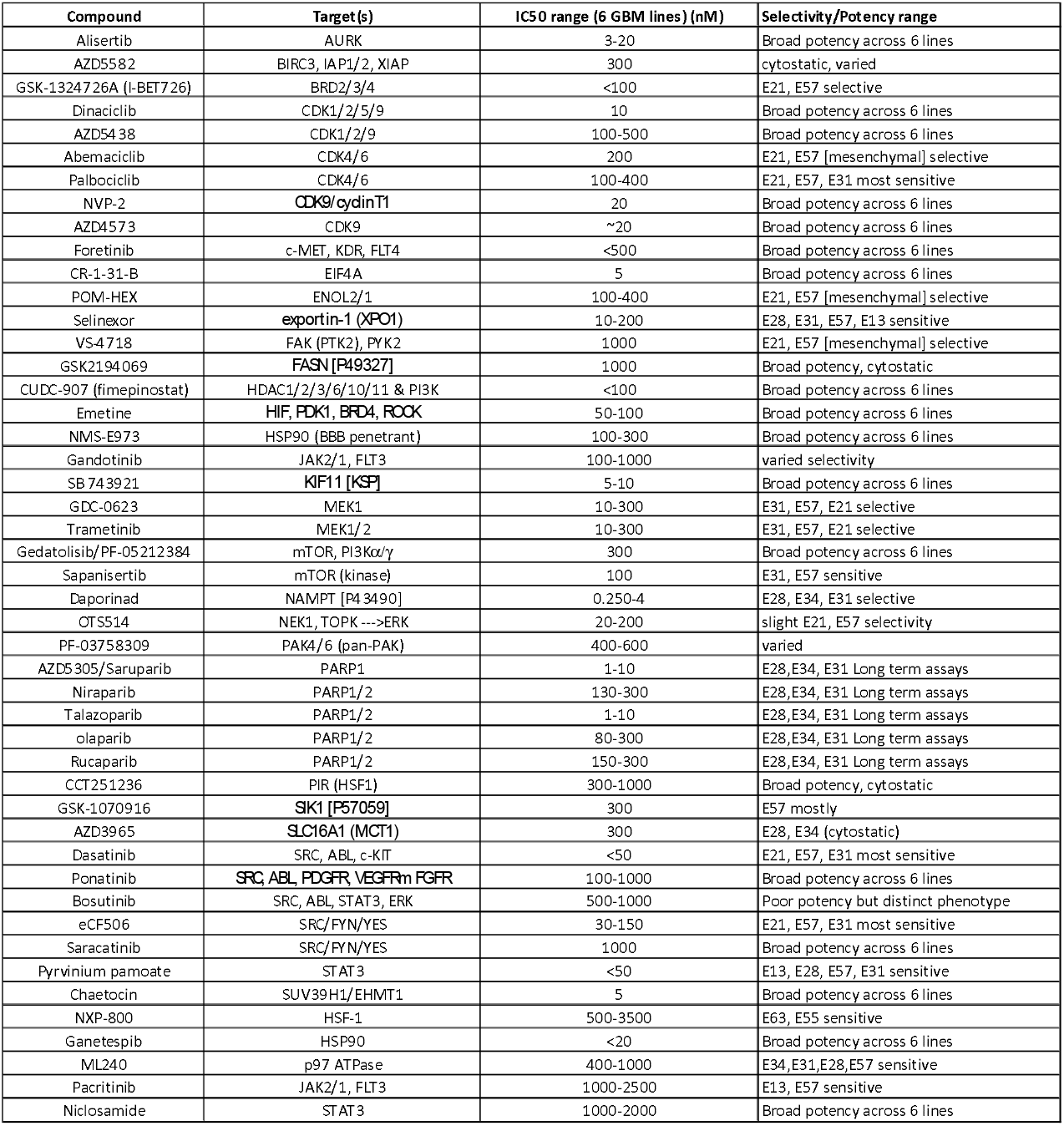
Validated compounds of primary interest. Compounds are listed in alphabetical order of their annotated primary target(s). The IC_50_ range (min to max) across 6 cell lines is given with indicated selectivity between lines (where applicable).

**Table 2:** Screening libraries and concentrations used.

| Library | # Compounds | Source | Concentrations Used (nM) | Controls (nM) |
| --- | --- | --- | --- | --- |
| TargetMol-Anti-Cancer | 330 | TargetMol #L2110 (2018) | 10000, 100, 10, 1nM | Paclitaxel (5nM),<br>Staurosporine (100 nM) |
| Kinase Chemogenomic Set (KCGS) | 187 | Structural Genomics Consortium<br>https://cancertools.org/the-kinase-chemogenomic-set-kcgs/ | 1000, 100 nM | Paclitaxel (5nM),<br>Staurosporine (100 nM) |
| Prestwick FDA | 1280 | Prestwick Chemical (~2016)<br>https://www.prestwickchemical.com/screening-libraries/prestwick-chemical-library/ | 10000, 1000 nM | Staurosporine (1000nM) |
| LOPAC<sup>®</sup> (Library of Pharmacologically Active Compounds) | 1280 | Merck/Sigma-Aldrich (LO4100) | 3000, 500 nM | Staurosporine (1000nM) |
| Comprehensive Anti-Cancer Compound Set C3L | 789 | Custom (reference 53; Athanasiadis et al, 2023) | 3000, 300, 30, 3 nM | Staurosporine (1000nM) |

### High Content Cell Painting Screen, Analysis & Validation

Our optimised GSC high content phenotypic screening and analysis workflow is outlined (Figure 1C) and described in detail in the material and methods section.

Briefly, cell lines were seeded (500-1500 cells per well) onto laminin pre-coated 384-well plates, followed by compound addition at 24 hours using automated liquid handling platforms. DMSO was used as a negative (compound vehicle) control (n = 48 wells per plate minimum) with staurosporine (1µM, n= 16) typically used as an assay landmark control for cell death. After 72 hours compound exposure, cells were subjected to live cell labelling of mitochondria with MitoTracker™, followed by fixation, permeabilization, and additional staining with the remaining Cell Painting dyes (see Materials & Methods). Images for phenotypic analysis were acquired using an ImageXpress Confocal high-content microscope with six fields of view (20× objective) over five channels, collecting 11,520 images per 384-well plate. Each compound library was screened separately across all six GCGR cells lines. In total 3,866 compounds were screened (varied doses), ∼2.2 million images collected (21 Tb) representing a dataset of >62,000 datapoints. Image analysis was carried out *via* a custom CellProfiler pipeline and secondary multiparametric data analysis was performed using StratoMineR™ (Core Life Analytics)[54] and TIBCO Spotfire® software (Revvity) - see methods. Representative principal component plots (PC1-3 of 20 principal components, covering >60% of explained variance) were created for LOPAC, Prestwick and C3L libraries (Figure 2, panels A, B, C & Supplementary Figure 4) to visualize distribution of active compounds from DMSO controls in phenotypic space. Further dimensionality reduction of these principal components was carried out, creating a phenotypic (Euclidean) distance for each compound, dose and cell line, relative to DMSO controls. The corresponding p-value to this phenotypic distance was plotted against cell survival (z-scores), based on DMSO normalised nuclei counts (Figure 2D, E & F). Standard high-throughput screening (HTS) quality control assessments were also carried out *via* Nuclei Count outputs from DMSO and staurosporine controls i.e., signal-to-noise *via* Z-prime robust (>0.35 minimum) across each plate (Supplementary Table 2). Image quality modules in CellProfiler were used to flag artefacts/blurred images, as well as additional compound triage *via* direct image assessment for putative hits.

**Figure 2.**
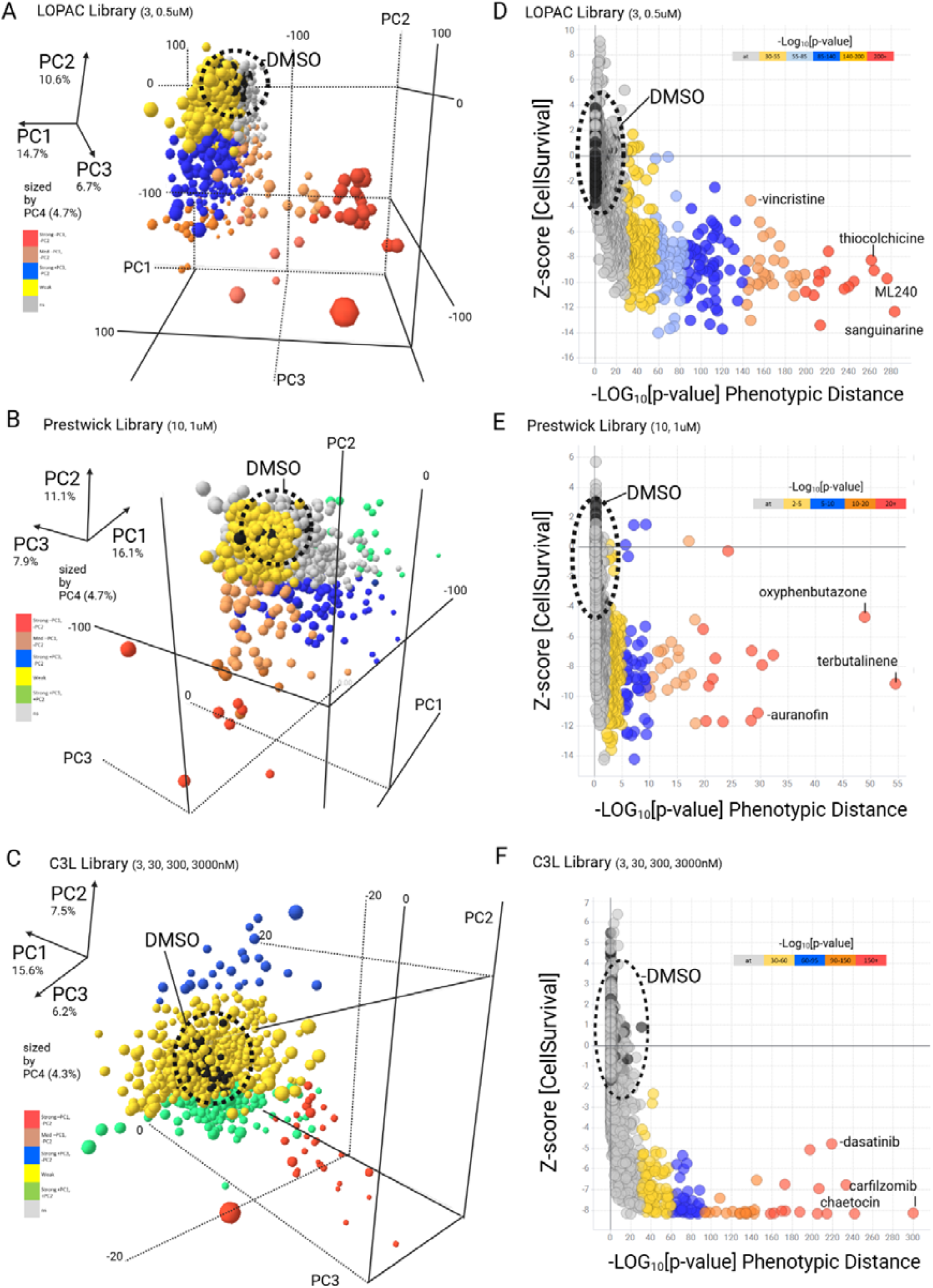
High-content image analysis/dimensionality reduction of drug screening data. A,B,C. Representative data of high content analysis across six GBM stem cell lines and three compound libraries (LOPAC, Prestwick FDA & C3L) by 3D Principal Component Analysis (PCA). DMSO controls circled in Black. Magnitude of vector co-ordinates for PC1,2,3 (sized by PC4) are indicated by colour: Red-pink (strong -PC1, -PC2), Blue (Strong +PC3, -PC2), Green (Strong +PC1, +PC2), Yellow/grey (weaker phenotypes). PC1-4 factor loadings are given (%). D,E,F. Phenotypic distance of screened compounds. We employed 20 principal components (>60% explained variance) to compute a phenotypic distance (Euclidean) for each compound[54]. The corresponding p-values are plotted against cell survival (normalised z-scores, Y-axis). Legend indicates strength of -Log_10_[p-value] distribution (5 bins, red = maximal effect, reducing to grey where at = arbitrary threshold). Several example hits are labelled.

We examined the degree of clustering of compounds with a phenotypic response far from the DMSO controls utilizing a combination of k-means clustering and silhouette score analyses (Supplementary Figure 5A,B)[55]. This analysis did not reveal a noticeable cluster structure and thus suggests a lack of morphological or phenotypic diversity of treatment effects across all screened libraries. Additionally, the distribution of the phenotypic distances across individual cell lines and libraries shows substantial heterogeneity in drug sensitivity, with particularly marked differences between the Prestwick and C3L libraries (Supplementary Figure 5C).

Initial hit identification was based on two phenotypic criteria, a cell survival threshold (Nuclei Count Z score <-3) and phenotypic distance threshold (negative Log_10_[p-value] >2) capturing changes in cell number and/or morphology following compound treatment in two or more GSC lines. Hit compounds were further triaged by removal of excessive examples of overtly cytotoxic agents (including common anti-cancer agents which have historically been investigated across pan-cancer indications, including GBM), such as microtubule disruptors, topoisomerase poisons and proteasome inhibitors. Compounds active at only high concentration (3µM) were de-prioritised. Short-listed hits were taken forward for further investigation.

Two hundred and eleven compounds (8.3% hit rate) were repurchased from different suppliers from that of the original screening libraries, for validation by dose response assay. Fresh compound stocks were prepared in 384-well format in a seven-point semi-log dose response (3-3000nM, 40 compounds per plate, with DMSO control wells n= 28). Fresh cell stocks were thawed and the validation protocol was identical to the primary screening assay, with the addition of a normal neuronal stem cell control line (NS69FB_B). Nuclei count data were also extracted from the phenotypic features and the validation set was analysed via both multiparametric and univariate analysis, with normalisation to DMSO wells (Figure 3A-C). The multiparametric analysis is presented as -Log_10_[p-value] Phenotypic Distance across all cell lines (Figure 3A) and we also converted dose response data to normalised Area Under the Curve (nAUC, scaled between 0-1 with 100% death = 0) for validation ranking (nAUC <0.85 in any line) to indicate the potential of achieving a therapeutic window (Figure 3B). Examples of nAUC derived dose response data across each validation plate is shown (Figure 3C) with comparative dose responses between 3 example compounds shown: alisertib (Aurora A kinase inhibitor), OTSSP167 (MELK inhibitor) and APIO-EE-07 (RSK1/MSK2 inhibitor) (Figure 3D). In spite of similar dose response profiles between alisertib and OTSSP167, these compounds occupy distinct phenotypic space (Figure 3E and representative images, Figure 3F), hence this approach allowed us to identify compounds with comparable cell survival, while inducing different phenotypic changes via distinct MOAs.

**Figure 3.**
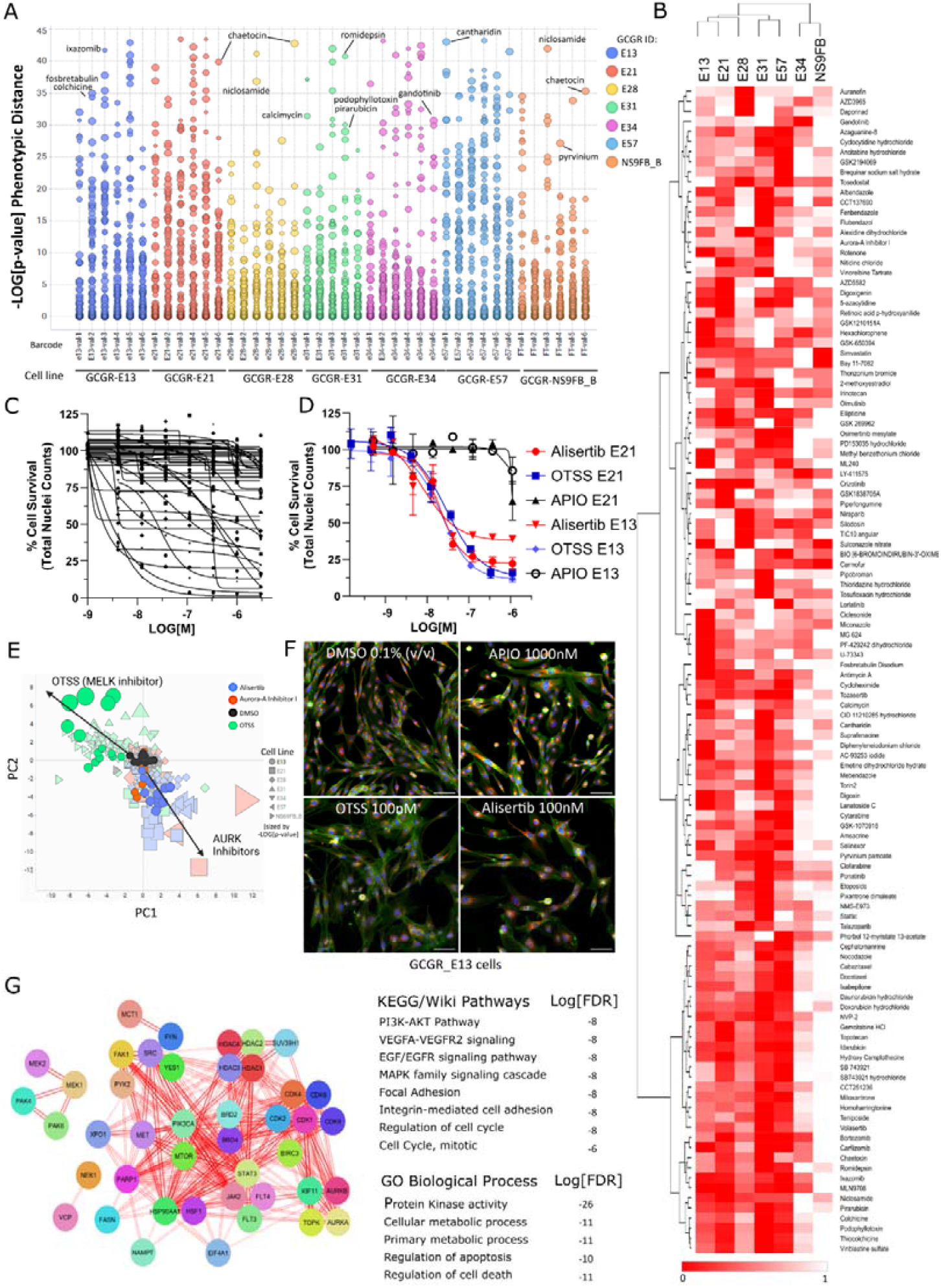
Hit validation from primary screening. A. Scatter plot representing phenotypic distance (negative-Log_10_[p-value] across 211 compounds and 6 GSC lines and 1 non-transformed NS cells (coloured by cell line, annotated by barcode and sized by dose (∼3nM - 1µM). Example hits are labelled. B. Representative heatmap of normalised Area Under the Curve (nAUC) data from dose response data (normalised nuclei counts) across 6 GSC lines and a normal neuronal stem cell control line (NS69FB_B), scaled between 0 (dead) and 1 (100% viable). Potency (low AUC) is indicated in red (relative shading by row). Hierarchical clustering by Euclidean distance/complete linkage. C. Representative normalised dose response data from a 384w plate (40 compounds per plate, 7 pt dose response, DMSO n=28) used to derive the nAUC. D. Example of dose response validation of alisertib (AURK inhibitor, red) vs OTSSP167 (MELK inhibitor, blue) and relatively inactive compound APIO-EE-07 (RKS/MSK2 inhibitor, black) across E13 and E21 cells. E. Example of distinct principal components/phenotypic space occupied by OTSSP167 (green) vs alisertib (blue) and Aurora A Kinase Inhibitor (red) over multiple doses vs DMSO (black). E13 cells are highlighted (circles) with vector co-ordinates indicated (black arrows). F. Representative images (E13 cells) displaying differing phenotypes between alisertib, OTSSP167 and APIO-EE-07 (vs DMSO) in spite of comparable IC_50_ curves. Staining: (Hoechst (blue), Phalloidin/WGA (green), Endoplasmic reticulum (red). Scale bar 100um. G. STRING-Cytoscape network & enrichment analysis of validated hits, by annotated target(s) search (sources: PubChem, Selleck Chemicals, ChEMBL, canSAR.ai). Lines indicate confidence of molecular interactions (edge score filter 0.4, text mining filter 0.06) with Gene Ontology (GO), KEGG and Wiki terms associated the enrichment analysis, with Log_10_[False Discovery Rate [FDR] is given. See also Supplementary Table 3.

In this study, validated compound data lists were generated, including separate phenotypic distance and cell survival assay endpoint lists (Table 1 & Supplementary Tables 1 & 2). One hundred and sixty four (164) compounds have been validated in dose response studies (Supplementary Table 1), including 143 compounds originally identified among the 211 hits in the primary screen (68% validation rate) and 21 alternative compounds, selected against target classes represented in the hit list. As expected, there are clear indications of in vitro sensitivity of GSCs to topoisomerase poisons, anti-metabolites, proteasome and HDAC inhibitors, as well as particular sensitivity to microtubule targeting agents (taxanes, etc). A prioritized short list (47 compounds) was generated to explore more novel molecular targeted therapies which displayed activity upon GBM and selectivity across a heterogenous GSC panel (Table 1). A network enrichment analysis of this prioritised list was carried out, using the annotated primary protein target(s) for each compound (sources: PubChem, Selleck Chemicals, ChEMBL, canSAR.ai databases), which demonstrates broad coverage of key target linked pathways, including PI3K-AKT, MAPK and EGFR signalling (Figure 3G and Supplementary Figure 6, Supplementary Table 3). Agents of interest involve those targeting cell adhesion (FAK, SRC, YES, FYN), transcription and translation (JAK2/STAT3, CDK9, HDAC, BRD), MAPK pathway/crosstalk (MEK1,2, PAK4,6), DNA damage/repair (TOPO, PARP), mitosis (AURK, KSP), metabolism (FASN, NAMPT) and protein folding/homeostasis (HSP90, HSF, VCP). There is clear chemical and target diversity within this validated hit list, beyond the expected highly potent cytotoxins, which provides a preliminary shortlist for rational drug-target selection and future systematic drug combination strategies. Any such approach would ideally be compatible with current therapy of post-surgical DNA damaging treatments.

### Secondary Assays/Exploration of target classes: Targeting Transcriptional Regulation

As it is beyond the scope of this article to follow up all of these hits in secondary assays, we have provided our full list of validated hit compounds for the GBM research community to explore and perform follow up investigation across similar and alternative GBM models. Within the scope of the current study, we have partially characterised two validated target classes (HDAC and cyclin dependent kinase inhibitors) which demonstrated highly potent (low nM) activity upon GSC survival (Supplementary Table 1).

Targeting transcriptional regulation (1): HDAC inhibitors. We further explored a focussed library of 54 structurally distinct HDAC inhibitors by dose-response, as our screening data demonstrated the significant potency of this class of compounds across our GSC panel (e.g., romidepsin, panobinostat) (Figure 4). Commercially available HDAC inhibitors were purchased and 384-well dose response plates were prepared and screened against the same six GSC line panel used for primary screening. Compounds were ranked by potency using the cell survival endpoint (normalised nuclei counts) (Figure 4A). Unsurprisingly, the most potent compounds were pan-HDAC or HDAC1/2 inhibitors, although the top ranked compound (with IC_50_ values below 20nM across all cell lines) was the dual HDAC/PI3K inhibitor, fimepinostat (CUDC-907) which is currently undergoing phase II clinical trials in paediatric brain tumours (NCT03893487). Fimepinostat, romidepsin and panobinostat were further validated by performing dose-response and cell cycle analysis (Figure 4B, Supplementary Figure 7A-C). Proliferation/cell survival over 72 hours is non-selectively inhibited across all cell lines, with cell cycle effects generally being limited to either small G_0_/G_1_ effects or strong G_2_/M arrest (Supplementary Figure 7A-C). Live cell imaging on E13 and E57 cells over 96 hours, with an intracellular caspase activity reagent (NucView-488, Biotium), showed a gradual but continuous induction of apoptosis from 24 hours (Figure 4C,D) with corresponding loss of proliferation from 24-36 hours (Figure 4E,F). Staurosporine (STS, 300nM) was used as a positive control for the apoptotic signal.

Targeting transcriptional regulation (2): CDK9 inhibitors. Cyclin dependent kinase inhibitors have been extensively investigated for their anti-cancer activity[56, 57] and several CDK inhibitors feature in our validated hit list, including approved CDK4/6 inhibitors abemaciclib and palbociclib. In particular, the CDK1/2/5/9 inhibitor, dinaciclib, as well as the selective CDK9 inhibitor, NVP-2[58], both show potent activity across multiple GSC lines (11-40 nM IC_50_ across the six GSC lines) (Figure 5A,B). In addition, a more potent, second generation CDK9 inhibitor, AZD4573, was included in this validation stage and also demonstrated low nM (7-27nM) activity across the 6 GSC lines (Figure 5A,B).

**Figure 4.**
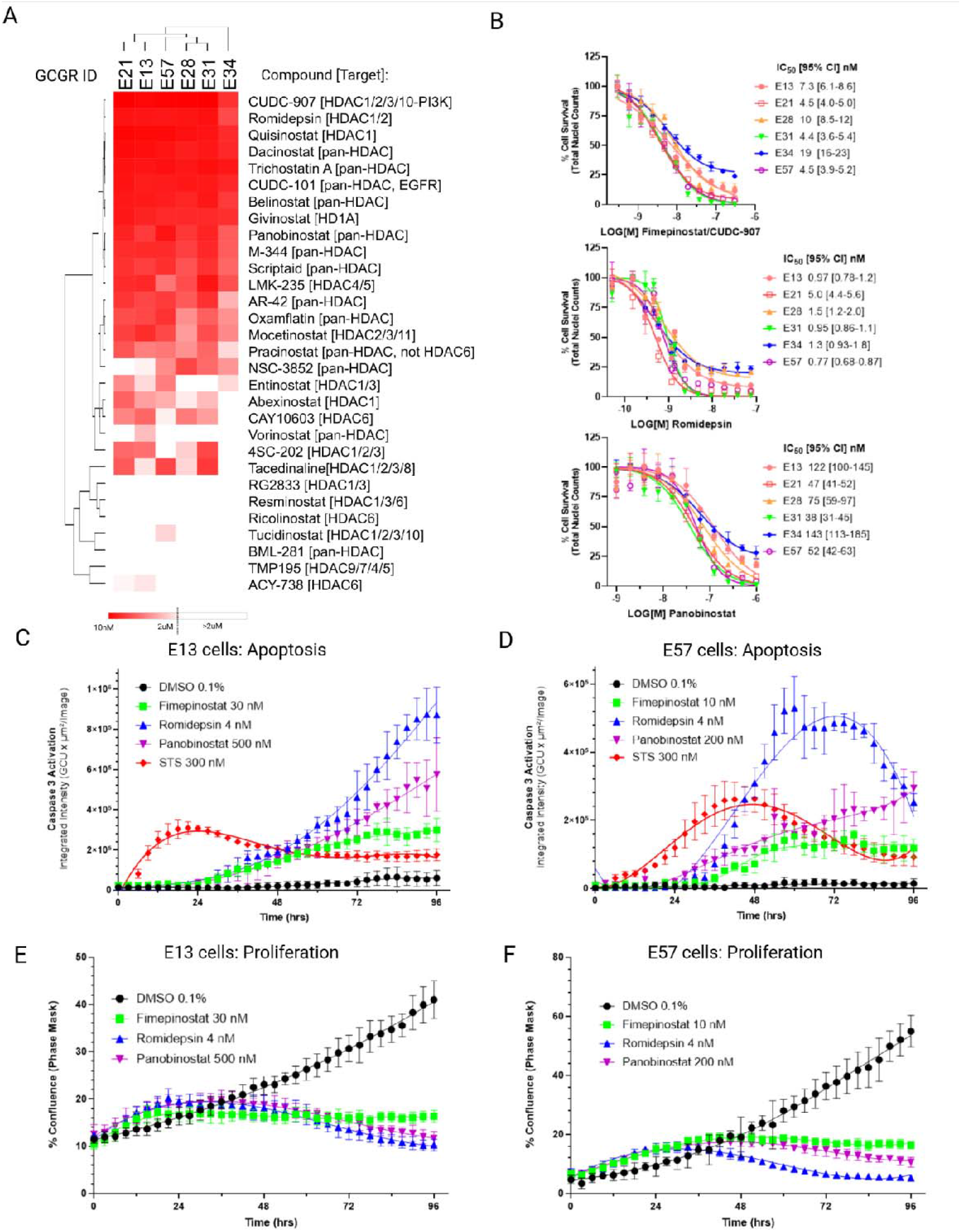
HDAC inhibitor library screening and profiling. A. Representative heatmap of IC_50_ values of HDAC inhibitors on GSC lines, quantified by DMSO normalised nuclei counts at 72 hours. The lowest IC_50_ compounds indicated in red (from 10nM to 2 µM, inactive >2µM indicated in white). Hierarchical clustering by complete linkage, Euclidean distance using Morpheus (https://software.broadinstitute.org/morpheus). B Dose response of fimepinostat (CUDC-907)), romidepsin & panobinostat with indicated IC_50_ values (nanomolar, nM) and 95% Confidence Interval (72 hours, n=3). C&D. Quantification of apoptosis by live cell imaging over time (imaged every 3 hours for 96 hours) at indicated IC_80_ values with staurosporine control (STS, 300nM, red). E & F. Quantification of proliferation over time (imaged at 3-hour intervals for 96 hours), at indicated IC_80_ values. DMSO control (0.1% v/v) is represented in black versus fimepinostat (30nM, green), romidepsin (4nM, blue), panobinostat (500nM, purple).

**Figure 5.**
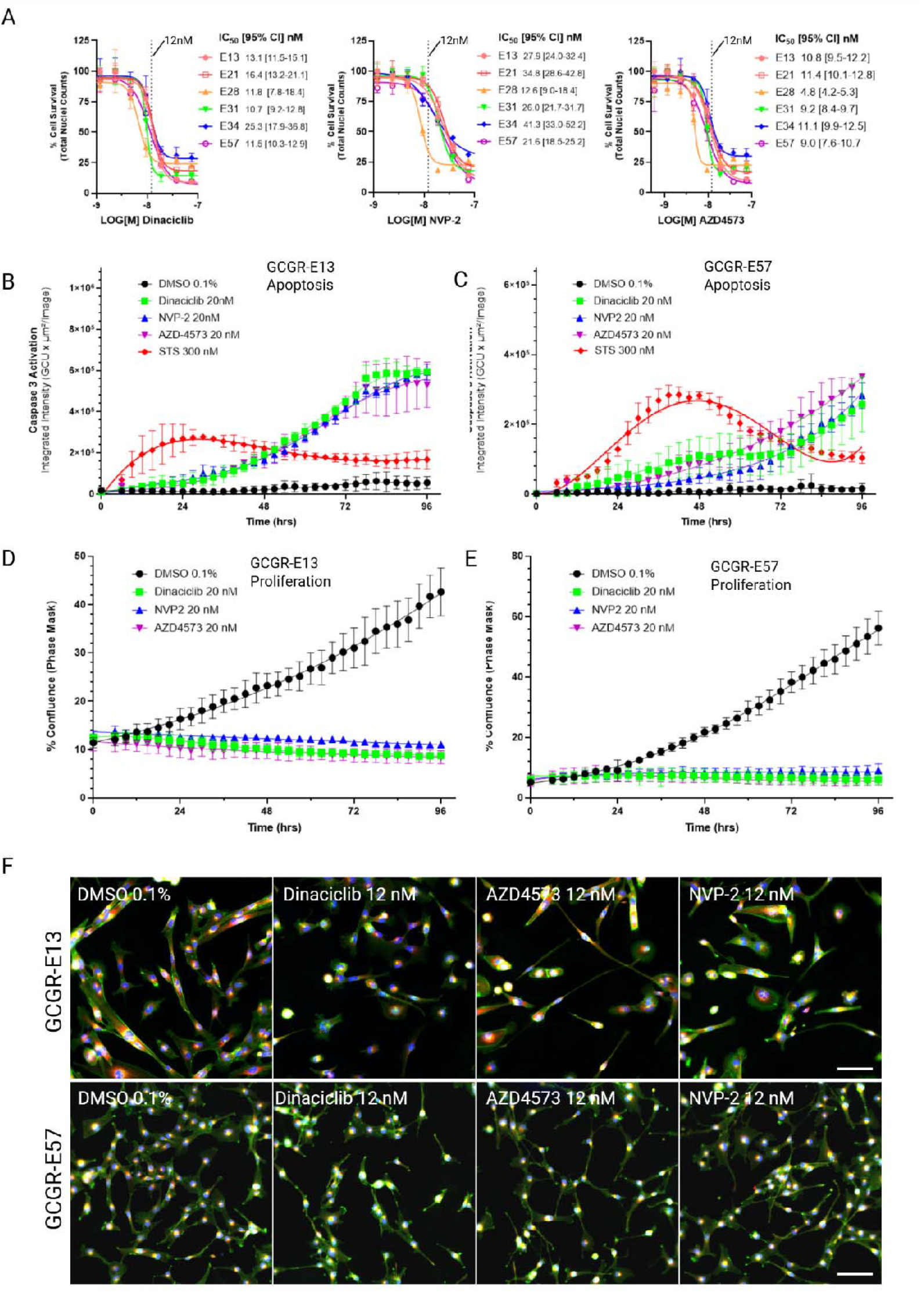
Targeting CDK9 in GBM. A. Dose response validation and IC_50_ values with 95% confidence interval (95% CI) of Dinaciclib, NVP-2 and AZD4573 (n=3). Twelve nanomolar (12nM) concentrations indicated with dotted line. B,C. Quantification of apoptosis by live cell imaging over time (imaged at 3 hour intervals for 96 hours), all compounds at 20nM concentrations with staurosporine control (STS, 300nM, red) and D,E. Quantification of proliferation over time. DMSO control (0.1% v/v) is represented in black, versus dinaciclib (green), NVP-2 (blue), AZD4573 (purple). F. Representative images of Dinaciclib, AZD4573 and NVP-2 at IC_50_ value (12nM) vs DMSO on E57 and E13 cells.

Dinaciclib is a multi-CDK inhibitor which has gained clinical interest in that it targets both cell cycle progression (via CDK1/2) and transcription (via CDK9), with this ‘dual targeting’ mode shown to be effective when combined with cisplatin in ovarian and endometrial cancers[59, 60]. Validation of dinaciclib, NVP-2 and AZD4573 across our GSC panel indicates broad potency at 72 hours across all the cell lines tested (Figure 5A), with G_2_/M arrest being initiated at concentrations in excess of 20nM ie below the IC_50_ values observed (Supplementary Figure 7D-F). Live cell imaging of these compounds, on E13 and E57 cells over 96 hours and in the presence of intracellular caspase activity reagent, showed rapid induction of apoptosis beginning at <24 hours from compound addition (Figure 5C,D) with corresponding immediate inhibition of proliferation, as observed from % confluence quantification (Figure 5E,F). Representative images of the phenotypic effects of these compounds at their IC_50_ concentration are shown (Figure 5G). This suggests transcriptional dysregulation via CDK9 inhibition is having a significant effect upon cell survival, independent of CDK1/2 inhibition. Overall, AZD4573 has improved cellular potency compared to dinaciclib & NVP-2 (Figure 5A,B,G: compare observable 12nM effects vs DMSO).

From these studies it is clear that targeting HDAC and CDK9, and hence transcriptional regulation, results in a cytotoxic benefit against GSCs, beyond the effects of prolonged cell cycle arrest (with the exception of the highly potent molecule, Romidepsin). To further explore the mechanism of action (MOA) of the CDK9 inhibitors on GSC lines, we performed NanoString™ transcriptomic analysis on E13 cells that were exposed to 20nM of Dinaciclib, AZD4573 and NVP-2, alongside DMSO controls, for 24 hours following compound addition. The extracted mRNA was quantified via the nCounter® system (Human Cancer Pathways, 784 genes including controls) and a differential analysis performed. After quality control filtration and removal of housekeeping/control gene sets (post normalisation), 467 genes were available for the differential expression & network pathway analysis, between untreated and treated cells. The results were further filtered by corrected p-value (Benjamini-Yekutieli FDR <0.05) and a minimum 3-fold change. Representative Volcano plots were generated for each compound with some highly significant genes labelled (Figure 6A).

**Figure 6.**
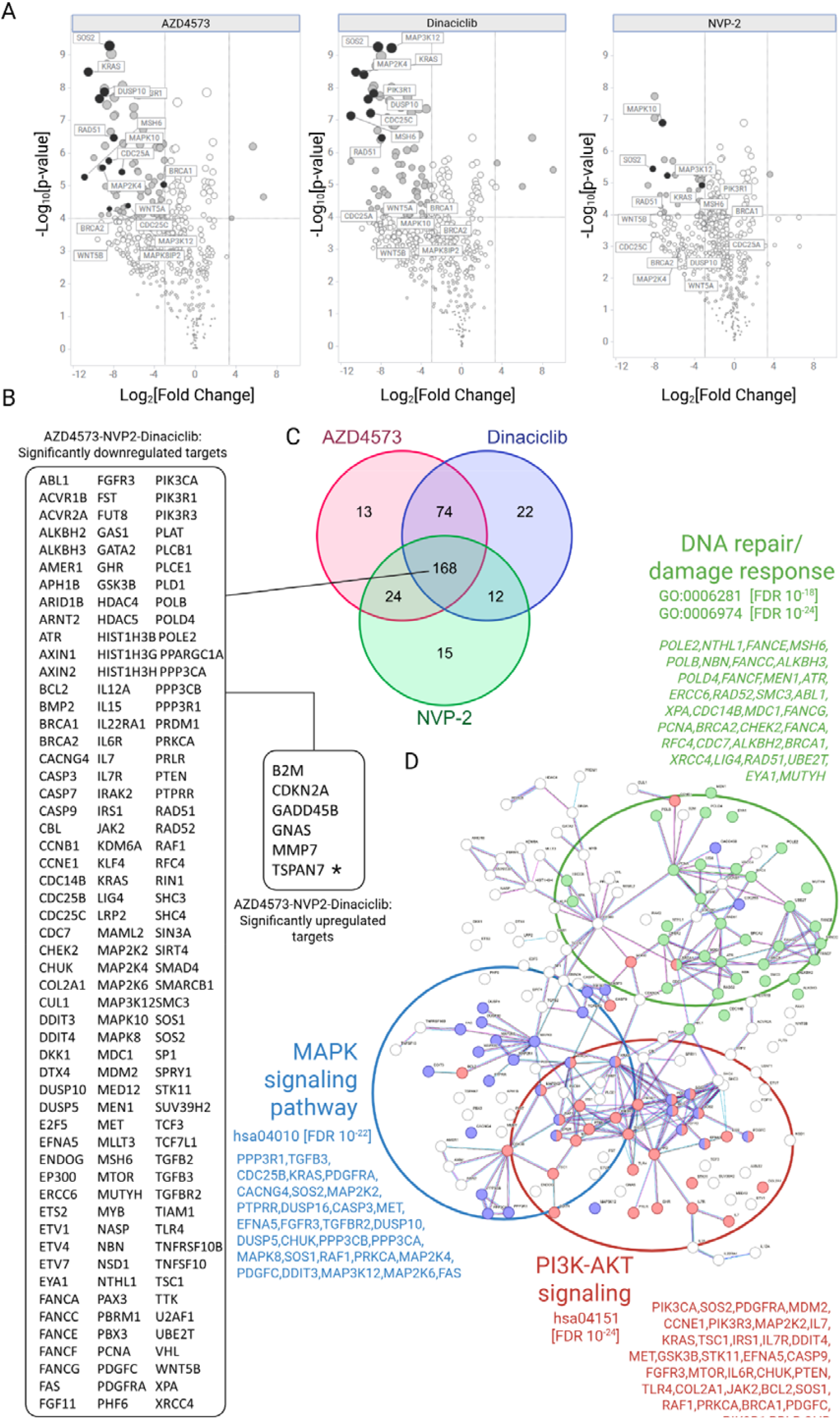
Differential gene expression (DE) analysis of CDK9 inhibitors on E13 cells from NanoString™ transcriptomics. A. Volcano plots of the differential analysis of E13 cells treated with AZD4573, dinaciclib and NVP-2 (20nM, 24 hrs), relative to DMSO control (0.1% (w/w)). B. Top 168 genes dysregulated by all three compounds (>3 fold change, corrected p-value <0.05). Large panel: downregulated genes, small panel: upregulated genes. TSPAN7 is highlighted with *. C. Venn diagram comparing gene hits across all three compounds. D. Network enrichment analysis (STRING) of top 168 genes. Enriched genes involved in DNA damage and repair (green), MAPK signalling (blue) and PI3K-AKT pathway (red) are highlighted. GO & KEGG terms with FDR is included. Enrichment tables and further data are provided (Supplementary Table 5, Supplementary Figure 8).

Not surprisingly, a large number of genes (168) were significantly dysregulated across all three compounds (Figure 6B), visualised by Venn diagram (Figure 6C). STRING network enrichment analysis (high confidence filter 0.700, no textmining) of these 168 genes (Figure 6D) showed significant enrichment of processes downregulating the MAPK cascade (e.g. KRAS, MAP2K4/MEK4, MAPK8/JNK1, MAP2K6/MEK6, DUSP10), PI3K-AKT signalling (e.g. PIK3CA/CB/CD [p100α,β,δ], IRS1, FGFR3, mTOR, MYB), DNA damage response/repair (e.g. RAD51, RAD52, BRCA2, PCNA, POLE2) and apoptotic signalling (Figure 6D. MCL-1, an anti-apoptotic BCL2 family member, is a reported CDK9 transcriptional and protein target[61]. This probe is not present in the NanoString™ panel, however, BCL2 and BCL2L1 mRNA were downregulated >10 fold, by all three compounds. Very few genes were significantly upregulated for each compound, but notably TSPAN7 (tetraspanin 7) was upregulated 5-8 fold – a protein where low expression is associated with poor prognosis in glioma and negatively correlates with immune infiltration of tumour associated macrophages[62]. The complete network analysis for each compound is provided (Supplementary Figure 8A-I) with enrichment tables and additional data (Supplementary Table 5).

Finally, we investigated potential synergistic drug combinations with some of our most potent hits which are currently clinically approved drugs: specifically, dinaciclib against romidepsin (pan-HDAC inhibitor), idarubicin (TOPOII inhibitor) and ixazomib (proteasome inhibitor) (Supplementary Figure 9A-O). Querying the Cancer Dependency Map database[63] (depmap.org/portal), a strong ‘cell death’ correlation was observed between dinaciclib and romidepsin/idarubcin/ixazomib (Supp Fig 9B,G,L, R^2^ = 0.62 to 0.67) across 564 human cancer cell lines (PRISM Repurposing Public 24Q2 dataset). In order to establish if this correlation could predict synergistic interactions, we carried out multiple drug combinations (7×7 dose response matrices) across E13 and E57 cells (72 hours exposure) with quantification from nuclei staining/counts. Whilst the drug combination of dinaciclib with either romidepsin, idarubicin & ixazomib does indicate highly potent additive effects upon GSC survival, synergistic activity was not observed in this instance (Supplementary Figure 9E,J,O).

## DISCUSSION

The clinical treatment of GBM has not significantly improved median patient survival in more than 20 years, with advances in the genomic characterisation of the disease only highlighting the complexity of the clinical challenge. To address this challenge a number of phenotypic screens of drug libraries and drug combinations performed in complex ex vivo or in vitro GBM models have recently been reported[64, 65]. However, screens reported to date have been limited in terms of the number of compounds, phenotypic endpoints and omission of defined tissue culture conditions which preserve GBM stem cell characteristics. In this study, our aim was to attempt to address the unmet clinical need in GBM by applying a comprehensive target agnostic, high content phenotypic screen, of thousands of compounds across a well characterized panel of patient-derived GBM stem cell models. The drug libraries were selected to support repurposing of existing oncology drug discovery programs and included Phase I passed or FDA/EMA approved small molecules and drugs with high target selectivity across known oncology targets. We have chosen to not exclusively utilise blood-brain-barrier (BBB)/CNS penetrant drug libraries in order to provide a comprehensive and unbiased evaluation of distinct drug and target classes upon GSC phenotypes. It is our hope that this will be informative towards future medicinal chemistry efforts in generating CNS penetrating small molecules against such targets, or possible incorporation of validated hit compounds into emerging blood-brain-barrier penetrating nanoparticles, designed to effectively deliver drugs and drug combinations to GBM [51-53]. In total 3,866 compounds were screened over multiple concentrations, in order to capture the most potent dose-dependent cellular phenotypes beyond simple viability measurements. Our GBM models comprised six heterogeneous, patient-derived GSC lines, representing three of the major transcriptomic subtypes (CLA, MES, PRO). We applied the multiparametric high content ‘Cell Painting’ assay to screen five drug libraries across the six GSC lines, and provide a validated list of 164 biologically active compounds on human GSC survival and phenotypic activity (Table 1 & Supplementary Tables 1 & 2).

Common cytotoxic agents, such as microtubule targeting agents, topoisomerase poisons, proteasome and HDAC inhibitors, are prevalent in our hit list and many have previously been investigated in GBM (and other cancers). Second generation taxanes, such as cabazitaxel, have shown modest effects with respect to mOS in phase I/II clinical trials (NCT01866449)[66]. The brain penetrant taxane, abeotaxane (TPI-287) showed some promise in phase I trials when combined with bevacizumab (NCT01933815), although confirmation of efficacy via phase II trials is required[67]. A recent meta-analysis of the use of the topoisomerase II inhibitor etoposide (44,850 patients over 624 studies published from 1976 to 2011) showed improved median OS (15.66 months vs 13.27, p=0.026) over topoisomerase I inhibitor, irinotecan, which actually showed reduced median OS (10.20 vs 13.55 months, p=0.008). Results are pending for a novel brain penetrant topoisomerase II inhibitor, berubicin (WP744), which has been granted Fast Track Designation by the US Food & Drug Administration[68]. Proteasome inhibitors have been clinically useful in treating blood cancers, such as myeloma[69], unfortunately a phase III study of the brain penetrant proteasome inhibitor, marizomib, did not improve mOS against unstratified patients[70]. The HDAC inhibitor class, such as romidepsin, in general have shown considerable promise as single agents for GBM treatment in the pre-clinical setting, but without translational success[71]. Clinical trials as combination therapy, for example combining panobinostat or vorinostat with radiation, bevacizumab and/or temozolomide have not proved particularly effective[72-74]. Arguably, this lack of clinical success could be attributed to trial design, limited predictive biomarkers, inadequate assessment of brain penetration, etc[75, 76]. Many of these agents have a narrow therapeutic window and significant systemic toxicities limiting their effectiveness. Nevertheless, novel drug delivery technology, such as antibody-cytotoxin conjugates, brain tumour specific nanoparticles[77-79] and intra-cranial drug delivery [80], could still render potent molecules clinically useful in the future.

Other molecular targeted compounds identified from our screen, include those targeting cellular adhesion, transcriptional regulation, DNA damage/repair, cell cycle/mitosis/proliferation, metabolism and protein folding/homeostasis, provide a basis for rational drug combination hypotheses and systematic co-inhibition strategies. For example, focal adhesion kinase (FAK) and SRC Family Kinases (SFKs) are downstream effectors of the integrin family of extracellular matrix receptors, and FAK is believed to influence GBM pathogenesis and brain tissue invasion[81-83] while SFKs are recognised drug targets in GBM[84-86] and PYK2 (FAK2)[87], Integrin-Linked Kinase (ILK)[88] and SRC all show high basal expression in our GSC models (Supplementary figure 2). Synergistic drug combinations of FAK or SRC inhibitors with MEK inhibitors have been reported in other cancers[89, 90] and, in a concurrent HTS project, we have found the combination of VS-4718 (FAK inhibitor) with trametinib (MEK1/2 inhibitor) to be quite effective in heterogeneous GBM in vivo models[91]. In fact, recently the combination of the dual RAF/MEK inhibitor Avutometinib with FAK inhibition has been shown to be effective against pre-clinical models of melanoma brain metastases[92]. Also, a phase I/II clinical trial combining avutometinib and defactinib (VS-6063) for NF1 ^del^ or BRAF^mut^ brain tumours has been initiated (5G-RUBY trial, NCT06630260) while the US Food & Drug Administration has recently granted accelerated approval for the avutometinib/defactinib combination for KRAS^mut^, low grade serous ovarian cancer (RAMP-201, NCT04625270, May 2025). Exploiting GSC vulnerabilities with respect to cellular adhesion and combining with other druggable routes (e.g. EGFR/MAPK/PI3K signalling) remains a viable strategy for GBM treatment. Additionally, we have identified distinct sensitivities to molecules targeting Heat Shock Factor-1 (HSF1; CCT251236, NXP-800), EIF4A (CR-1-31-B), SUV39H1 (chaetocin), FASN (GSK2194069), NAMPT (daporinad), Enolase-2 (POM-HEX), Exportin-1 (selinexor) and JAK2 (gandotinib, pacritinib) (Table 1). Furthermore, our examination of the basal expression of our GSC lines by reverse phase protein and cytokine array also reveals actionable targets via high expression of Forkhead Box A1 (FOXA1) and interleukin-8 (IL-8) (Supplementary Figure 2). FOXA1 is usually associated with hormone dependent cancers and can drive endocrine resistance in oestrogen-positive breast cancer[93], or MAPK activation in prostate cancer[94], in an IL-8 dependent manner. Targeting IL-8 via humanised anti-IL8 antibody[46] is potentially another combination strategy that could be systematically investigated alongside our prioritised hits (Table 1, Supplementary Table 1).

In this article we followed up two drug target classes, HDAC and CDK inhibitors which displayed low nanomolar activity upon inhibiting cell survival across the heterogeneous panel of GSCs. To further explore the HDAC inhibitor class we screened a collection of 54 HDAC inhibitors. Fimepinostat (CUDC-907) was outstanding as the most potent molecule, with <20nM IC_50_ values across the GSC panel. Fimepinostat, a dual PI3K/HDAC inhibitor, is currently undergoing clinical trials for paediatric/young adult brain cancers/solid tumours (NCT03893487), and is potentially a candidate for targeted drug delivery and drug combination strategies. We compared fimepinostat with structurally distinct HDAC inhibitors, romidepsin and panobinostat. While there was a varied cell cycle response between cell lines (Supplementary Figure 7), there was a common intra-cell line response between molecules, suggesting activity is most likely mediated through the anti-HDAC activity of fimepinostat rather than its anti-PI3K activity. Indeed, comparison between fimepinostat and potent, selective PI3K inhibitors (all tested at 30nM) demonstrate substantially weaker activity of PI3K inhibition upon GSC proliferation, with only dual PI3K/mTOR inhibitor, gedatolisib (PF-05212384), showing any significant effect (Supplementary Figure 10). While the lack of success in targeting GBM with PI3K monotherapy may indicate a lack of true oncogenic addiction to PI3K signalling, drug combinations with PI3K/mTOR/AKT inhibitors are still viable options against PTEN deficient, GBM tumours[95].

Targeting transcriptional regulation has previously been shown to be effective against cancer stem cells (including brain cancers)[96-98], hence we also characterised potent CDK9 inhibitors dinaciclib, NVP-2 (both screening hits) and a more recent selective CDK9 inhibitor, AZD4573. CDK9 is an atypical CDK in that it is a key regulator of RNA Polymerase II (Pol II) transcription initiation, elongation and termination, playing a critical role in transcriptional regulation[99]. Dinaciclib has near equal potency against CDK1/2 (& 5), whereas NVP-2 and AZD4573 are classed as CDK9 selective[100, 101]. AZD4573 has progressed to clinical trials (NCT04630756) and is well tolerated, although CNS penetrance and solid tumour efficacy remains to be determined. Radiosensitisation of cancer cells with CDK9 inhibitors and drug combinations with DNA damaging agents have previously been reported[102-104], including opportunities in GBM[105]. In our in vitro assays, dinaciclib, NVP-2 and AZD4573 are highly potent across all GSC lines tested, being strongly antiproliferative with predominant G_2_/M arrest and induction of apoptosis. Differential gene expression analysis of GSC cells exposed to CDK9 inhibitors, reveals highly significant decreases in the expression of genes regulating the cell cycle, DNA damage and repair, anti-apoptosis, PI3K and MAPK signalling (Figure 6, Supplementary Figure 8, Supplementary Table 5). It is noteworthy that, of the very few genes upregulated in response to CDK9 inhibition, Tetraspanin-7 (TSPAN7/TM4SF2) expression is increased 5-8 fold (Figure 6B). An increase in TSPAN7 expression in GBM is associated with improved prognosis and immunosuppression of the tumour microenvironment[62], potentially impacting GBM response to anti-PD1 or anti-PD-L1 immunotherapy when also combined with CDK9 inhibition. These results indicate that the CDK inhibitors are reprogramming the GSCs into a more vulnerable state, leading to inhibition of cell survival and, together with previous publications implicating CDK9 as a therapeutic target in GBM[105], suggest targeting CDK9 is worth further exploration in other GBM models and with other drug combinations.

Overall, our screening results presented here serve as a foundation for a systematic approach to the identification, validation and prioritisation of novel therapeutic strategies in GBM including future drug combinations, that could be used concurrently or following standard treatment protocols for GBM. Importantly, we have used patient-derived GSC models, grown under stem-like conditions on a laminin rich ECM scaffold to prioritize follow up investigations of highly potent small molecules active across a genetically distinct and heterogeneous GSC panel. The objective of our phenotypic screening assay was to identify multiple drug targets and mechanisms-of-action which target heterogeneous GSC phenotypes and thus complements the existing focus on known oncogenic drivers that represent established drug target classes. We anticipate that this approach will identify new therapeutic targets and drug combination hypotheses that target GSC phenotypes rather than individual pathways and thus may be more resilient across heterogeneous patient populations and adaptive drug resistance mechanisms.

In conclusion, our high content screening followed by multi-cell line validation across primary and secondary phenotypic assays provides the largest unbiased survey of small molecule therapeutic classes and oncology targets upon GBM stem cell phenotypes that we are aware of. We provide our full list of validated hits active upon GBM stem cell morphology and cell survival for the research community to explore further. Future studies include expanded screening across larger target-annotated and diverse chemical libraries to identify novel chemical starting points and novel targets for new GBM drug discovery programs. In addition, further profiling of phenotypic screening hits across a larger number of GBM stem cells lines and integration of phenotypic with molecular data, in order to identify synthetic lethality relationships and/or predictive biomarkers of response, will support additional personalized medicine strategies for both existing and new drug target classes in GBM. Importantly this unbiased approach can identify new therapeutic opportunities for the majority of GBM patients which do not fall into well-defined subtypes characterized by well-known druggable driver genes thus expanding the scope and inclusion criteria of personalized medicine strategies in GBM.

## Supporting information

supplementary TablesS5

## ACKNOWLEDGEMENTS

This work was funded by a joint Cancer Research UK (C42454/A28596) and The Brain Tumour Charity award (GN-000676) to D.E., N.O.C. and M.C.F. G.M.M and the Glioma Cellular Genetics Resource were supported by the Cancer Research UK (CRUK) Centre Accelerator Award (A21922).

## Materials and Methods

### Cell culture

Cell lines were obtained from the Glioma Cellular Genetics Resource, Edinburgh (, https://github.com/GCGR) and cultured according to optimised conditions for glioma stem cell growth (gcgr.org.uk). Briefly, cells were cultured in DMEM/HAMS-F12 media (Merck, D8437) with B27, N2 supplements (GIBCO 17504-044, 17502-048), EGF/FGF (10ng/mL, Peprotech 315-09, 100-18b)), glucose (8mM, Merck, G8644), MEM non-essential amin acids (5mL, GIBCO 11140-035, 100X), BSA (0.012%, GIBCO 15260-037), β-mercaptoethanol (0.1mM, GIBCO 31350-010) and laminin (4-10µg/mL depending on the cell line, Cultrex 3446-005-01). Cells were passaged at ∼80% confluence (4-7 days) and dissociated using Accutase (Merck, A6964). Doubling times are approximately 60 hours (except E57 cells, doubling time ∼24 hours) with generally low passage ratios (1:3 to 1:6) to maintain no less than 20% confluence during cell expansion. Cell lines were submitted for STR profiling (ECACC, case number 21579, using a Promega Powerplex 16 HS kit) and were routinely monitored for mycoplasma infection (MycoAlert®, Lonza). All cell lines were expanded and banked in liquid nitrogen (∼24 vials each) for continuous turnover of low passage lines during screening.

### Compound screening

384-well plates (Greiner Bio-One, µClear, 781091) were precoated in growth media containing 10ug/mL laminin (20µL/well) and incubated for 2 hours at 37 ^°^C, 5% CO_2_ (minimum). Single cell suspensions in growth media were prepared (final seeding densities indicated in Supp Figure 1B) and added to the pre-coated plates (30µl/well, final laminin 4µg/mL). The plates were then incubated for ∼20 hours and inspected briefly, prior to compound addition using a Biomek automated liquid handler. Compound library plates (as assay ready daughter plates in 100% dimethylsulfoxide (DMSO), 10mM-1mM stocks, prepared via BioAscent compound management (bioascent.com)) were diluted with media to intermediate plates (1:50, 2% DMSO (v/v)) prior to addition to cells (1:20, final 0.1% DMSO (v/v)). Where required, the intermediate drug plates were further diluted to create reduced dose intermediate plates (with 2% DMSO (v/v)) prior to addition to cells (1:20, final 0.1% DMSO (v/v)). Negative controls (DMSO, 0.1% (v/v), n= 32-48 per plate) and positive controls (staurosporine, 1µM final conc., n= 16) for cell death were added to the intermediate compound plates (up to 320 compounds per plate). After the addition of compounds, the plates were then incubated at 37 ° C, 5% CO_2_ for 72 hours. Compound resupply details are provided in Supplementary Table 4.

### Cell painting

Live mitochondrial staining of cells was carried out by the addition of MitoTracker Deep-Red solution (3µM, 10X conc. in media at 5µL/well) using the Integra Viafill automated dispenser (Intergra Biosciences). The plates were then incubated for 30 minutes at 37 ^°^C, 5% CO_2_ followed by plate cooling and fixation by the addition of 15% formaldehyde in phosphate buffered saline (PBS) (20uL/well – final 4% formaldehyde in 75µL volume) using a multidrop combi reagent dispenser. After incubation for 20 minutes, the plates were washed (BioTek plate washer, Agilent, www.biotek.uk.com) with PBS and cell painting reagents prepared in 1% BSA/0.01% TX-100 in PBS (Table 3). After washing, PBS was removed and cell painting solution was added using a multidrop combi (20µL/well) followed by incubation at room temperature for 30 minutes. The staining solution was then removed, the plates washed with PBS, sealed (Starseal foil seals, Starlab, E2796-9792) then imaged immediately.

**Table 3.**
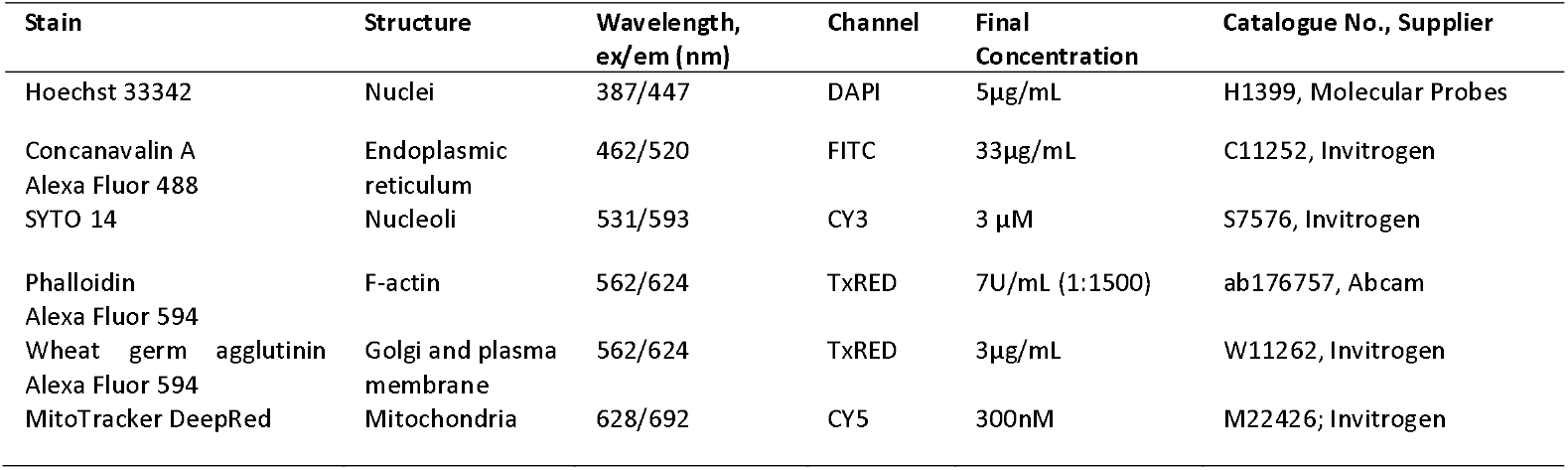
Cell painting reagents listed by target organelle, channel, final concentration & supplier details.

### Image acquisition

Images were acquired using ImageXpress Confocal HT.ai high content microscope (Molecular Devices LLC, version 6.5.0) with automated plate handling (GX series, Peak Analysis and Automation). Exposure settings for each channel were optimised and set for screening. Each well was imaged using a 20× objective and 6 fields of view (30 images per well over 5 channels, ∼33% well coverage, typically 75-150 cells per image)

### Image analysis, quantification and data processing/plotting

CellProfiler v3.1.5 image analysis software (cellprofiler.org) was used to create a custom analysis pipeline across 5 channels capturing 1155 features per cell (processed features & factor loadings/explained variance is provided in Supplementary Table 5). Images were analysed via parallel processing using a Unix/Linux high performance computing cluster (University of Edinburgh) and concatenated outputs included median aggregated, well level data. Each completed dataset (library across 6x GCGR lines) was then batch processed using StratoMineR high content analysis platform (stratominer.com). Briefly, stepwise processing in StratoMineR included metadata integration, removal of redundant features (correlation cut-off 0.99, typically 800-850 features remaining), QC and control annotation/flags, normalisation (to DMSO and sample wells, on a plate-by-plate basis), transformation (Skewness significance 0.0001), feature scaling (robust z score, by plate) and dimensionality reduction (principal component analysis [generalised weighted least squares], Oblique rotation, Ten Berge factor scores method, auto correlation cut-off with ≥20 factors chosen [model based on samples] to represent >60% of data variance [Scree plots]) and computation of phenotypic distance/p-values from principal component vectors[54].

Processed data was then exported and analysed further using TIBCO Spotfire analysis software (spotfire.com), including plot generation. Hierarchical clustering of normalised datasets was carried out using Morpheus (https://software.broadinstitute.org/morpheus) by complete linkage, Euclidean distance). Network and enrichment analysis of top ∼50 [supplier] annotated compound targets was carried out using STRING[106] & CytoScape (cytoscape.org)[107]. Cell cycle analysis was carried out from DNA content/nuclei images (Hoechst stain) using MetaXpress software (Molecular Devices LLC, version 6.5.0). Dose response data and curve fitting (non-linear regression) was carried out using GraphPad Prism (version 7.05). Synergy plots were generated using SynergyFinder+ web application[108] (https://synergyfinder.org/)

### Clustering Analysis and distribution plots

To quantify the phenotypic diversity of the screened compounds with z-score[cell survival] < -3 and - Log_10_[p-value] >2 we performed k-means clustering of the unique 245 compounds that passed this threshold using the principal components derived from the morphological features (20 PCs for libraries LOPAC and Prestwick; 10 PCs for C3L library). The degree of clustering was quantified by the k-means score defined as the within-cluster sums of point-to-centroid distances, summed across all clusters. The quality of the k-means clusters was determined with the silhouette coefficient S averaged across the unique compounds per library[55]. The silhouette coefficient varies between ™1 and 1, with S⍰=⍰1 indicating that compounds are in well separated clusters, S⍰=⍰0 indicating overlapping clusters, and S⍰=⍰™1 indicating incorrect assignment of clusters. The distribution of the variable -Log_10_[p-value]. Phenotypic Distance was calculated using violin plots per cell per library. These measures were calculated using R version 2023.9.0.463 with libraries base stats (function kmeans), cluster_2.1.4 (function silhouette) and ggplot2_3.4.3 (function geom_violin).

### Live Cell Imaging/Apoptosis Assays

GCGR-E13 (1000 cells/well) and GCGR-E57 cells (500 cells/well) were seeded (45µL/well) onto 384-well plates (Greiner uClear, 781910) pre-coated with 10µg/mL Laminin ECM. After overnight incubation, a solution of NucView-488 reagent (10X conc., 50µM) was added using an Integra ViaFill Liquid dispenser (5µL/well). The compounds were then immediately added using a D300e nanodispenser with back-filling of DMSO to 0.1% (v/v) for all wells, triplicate wells per dose (3 concentrations per compound, based on individual IC50 range). Staurosporine (300nM, 6 replicates per cell line) was used as a positive control for apoptosis induction against DMSO controls (0.1%, 6 replicates per cell line). The plate was then loaded into an IncuCyte S5 Live Cell imager with scans set at 3-hour intervals for 96 hours (phase and ‘green’ channels, 10× objective, 1 site). Image processing was carried using IncuCyte in-built analysis modules, with phase area defined as % Confluence/image from the first scan (∼t = 0+30mins). Apoptosis was quantified from images showing increasing Integrated Intensity from the NucView-488 emission signal (GCU x µm^2^ /Image).

### NanoString™ sequencing/differential analysis

mRNA was extracted from E13 cells (∼3×10^6^ cells each condition) after 24 hours treatment with AZD473, Dinaciclib, NVP-2 (all 20nM) & DMSO (0.1% v/v) in triplicate 10cm dishes, using RNeasy Plus miniprep extraction kit (Qiagen, #74134). Purified mRNA was quantified and diluted to 20ng/uL and submitted for NanoString™ nCounter® analysis against the Human Cancer Pathways Panel (NS_CancerPath_C2535). Samples were hybridised and immediately processed using the nCounter® Prep Station and Digital Analyzer (High Sensitivity protocol). QC checks utilised GeNorm to determine most stable reference genes and no QC Flags were raised overall. Differential analysis was carried out on normalised data (normalised to ∼20 housekeeping genes) using nSolver™ Analysis Software (version 4.0.70) following standard analysis pipelines. Exported data was replotted using TIBCO Spotfire software and also further analysed by hierarchical clustering (https://software.broadinstitute.org/morpheus) and STRING network analysis (string-db.org)[106]. Venn diagrams were generated from Molbiotools (molbiotools.com/listcompare.php)

### Forward-phase protein array: basal cytokine analysis

Conditioned medium was collected after 72 hours incubation. Microarrays were generated using an in-house Aushon BioSystems 2470 array printing platform. The arrays were blocked for 1 hour with Superblock T20 Blocking Buffer (Grace Bio Labs) at room temperature. Conditioned media from glioma stem cell samples were centrifuged at 1000× g for 5 minutes at 4 °C. Supernatants were collected and added to microarrays followed by incubation overnight at 4 °C. The arrays were washed three times for 5 minutes in PBS-T (PBS-Tween 0.1% v/v) and blocked for 10 minutes with Super G Blocking Buffer at room temperature on a rocking platform, then washed again washed three times for 5 minutes in PBS-T. Detection antibody mixtures (1:500 antibody dilutions in 5% bovine serum albumin/phosphate buffered saline tween-20 0.1% (BSA/PBS-T), 10% Super G Blocking Buffer) were made in microplates. Microarrays were clamped and 50 µL of each antibody was added to corresponding microarray wells. Microarrays were incubated for 1 hour on a rocking platform. Clamps were removed and microarrays were washed three times for 5 minutes in PBS-T. Microarrays were then blocked for 10min with Super G Blocking Buffer at room temperature and again washed three times for 5 minutes in PBS-T. Three millilitres (3 mL) of IRDye 800CW Streptavidin (LI-COR Biosciences) was diluted 1:5000 in PBS-T supplemented with 5% BSA, 10% SuperG Blocking Buffer. Microarrays were covered and incubated with IRDye on a rocker at room temperature for 30 minutes, then washed for 5 minutes, three times in PBS-T followed by three 5-minute PBS washes and finally washed with distilled water. Microarrays were dried then scanned at 795nm on the InnoScan 710 high-resolution microarray scanner (Innopsys Life Sciences). The data was normalised for protein concentration and background fluorescence via Microsoft Excel templates.

### Reverse-Phase Protein Array (RPPA): Basal analysis

GSC lines were cultured both as mono-layers on laminin-coated cell culture plates and as spheroid cultures on poly-hema coated plates. Cells were set up as mono-layer and spheroid cultures in parallel by seeding equivalent cell numbers from a parental flask with cells in log-phase growth. Cultures were grown until the mono-layer cultures were approximately 70% confluent and then both adherent and suspension cultures were harvested for lysis and preparation for RPPA. For Laminin coating, plates were incubated with Laminin at 10ug/ml for 1 hour and then washed with PBS prior to addition of cell lines. Culture media did not contain Laminin and the same media was used for adherent and suspension cultures. For Poly-hema coating plastic 2.4g of poly-hema was dissolved in 200ml of 95% Ethanol and heated to dissolve. The solution was centrifuged at 2500RPM to remove undissolved particles. 4ml was added per 10cm plate (ensuring the plate was completely coated). Plates were left open in a TC hood to air-dry and were rinsed x3 with PBS prior to addition of cells.

Mono-layer cultures were washed twice with PBS after removal of media and snap frozen on dry-ice and stored at -80 ° C. Spheroid cultures were pelleted by centrifugation and washed twice with PBS and then snap frozen and stored until lysis. Cultures were lysed in RIPA buffer supplemented with ROCHE PhosStop and cOmplete Mini inhibitor tablets (Merck, 04906845001 & 11836153001). Lysates were needle syringed five times to ensure complete lysis of cells. Protein quantification was carried out using Coomassie-plus assay (ThermoFisher #23228) and protein concentration was adjusted by to 1 mg/ml. Samples were prepared for printing by addition of 4x Sample buffer and heat denaturation at 95 ° C for 5 minutes.

All samples were diluted to 0.75mg/ml (D1), 0.375mg/ml (D2), 0.1875mg/ml (D3) and 0.09375mg/ml (D4) using PBS containing 10% Glycerol. Array spotting was carried out with the Quanterix 2470 Arrayer platform using 185μM pins. Each sample was spotted at four dilutions (D1-D4) onto single pad ONCYTE® SuperNOVA nitrocellulose slides at a 500μM spot-to-spot distance. 268 signalling pathway markers (and appropriate controls) were profiled in a standard RPPA assay as described briefly below. After spotting, the slides were incubated with antigen retrieval solution (1x Reblot strong) for 10 mins before being placed in a microfluidic structure to individually address the arrays with primary or secondary antibody solutions. Following blocking buffer (Superblock T20) for 10 mins the detection of marker antibodies was performed in a two-step sequential assay (1) incubation of an array with primary analyte-specific antibody in blocking buffer for 60 mins at room temperature and (2) removal of excess antibody by washing arrays with PBS-T, followed by a further incubation with blocking buffer and PBS-T washes, and incubation secondary antibody (Dylight-800-labeled anti-species antibodies diluted 1:2500 in Superblock T20) for 30 mins. After further washing and slide drying, the arrays were imaged in the Innopsys Innoscan 710 scanner. Blank signals were determined by omitting the primary antibody from step 1 and instead incubating the array with Superblock T20 alone, followed by step 2. Sample loading on arrays (for normalisation) was determined by staining one slide with fast green protein dye and scanning at 795nm. Microarray images are analyzed using Mapix software (Innopsys). The feature (spot) diameter of the grid was set to 270μm. The average signal intensity is determined for each individual feature and the median background from the adjacent area is subtracted from each feature signal leading to a net signal per feature. Data analysis is performed in a standard way: Fluorescence intensity for each feature on the array is measured. A test is performed for linear fit through the 4 point dilution series for all samples on all arrays using a flag system where R2 > 0.9 (green flag) is deemed good, >0.8 (amber flag) is deemed acceptable and <0.8 (red flag) is poor and may be excluded from data analysis. The median values from the 4 point dilution series are calculated and used as a measure of fluorescence intensity. Data is quantified as RFI (relative fluorescence intensity) values relating to relative abundance of total and phosphorylated proteins across the sample set.

