## Supplementary data and table 1 for "A comprehensive pharmacological survey across heterogeneous patient-derived GBM stem cell models"

**Supplementary Figure 1. Optimisation of seeding densities and incubation** A. Cell Line Molecular data (Glioma Cellular Genetics Resource (<https://github.com/GCGR>). B. Representative phase contrast images of GCGR cells at 72 hrs. GCGR-E13 (classical subtype, 1000 cells/well), GCGR-E28 (classical, 1500 cells/well), GCGR-E21 (mesenchymal, 1000 cells/well), GCGR-E57 (mesenchymal, 500 cells/well), GCGR-E31 (proneural, 1000 cells/well), GCGR-E34 (proneural, 1000 cells/well). Scale bar 400µM. C. Quantification of live cell imaging and growth curves for 384w optimisation. Doubling times ~60 hrs (except E57 cells ~24 hrs). D Optimisation of seeding density and quality control based on coefficient of variation (%CV) <20% (based on ‘Average Number of Nuclei’ per site/over 6 site).

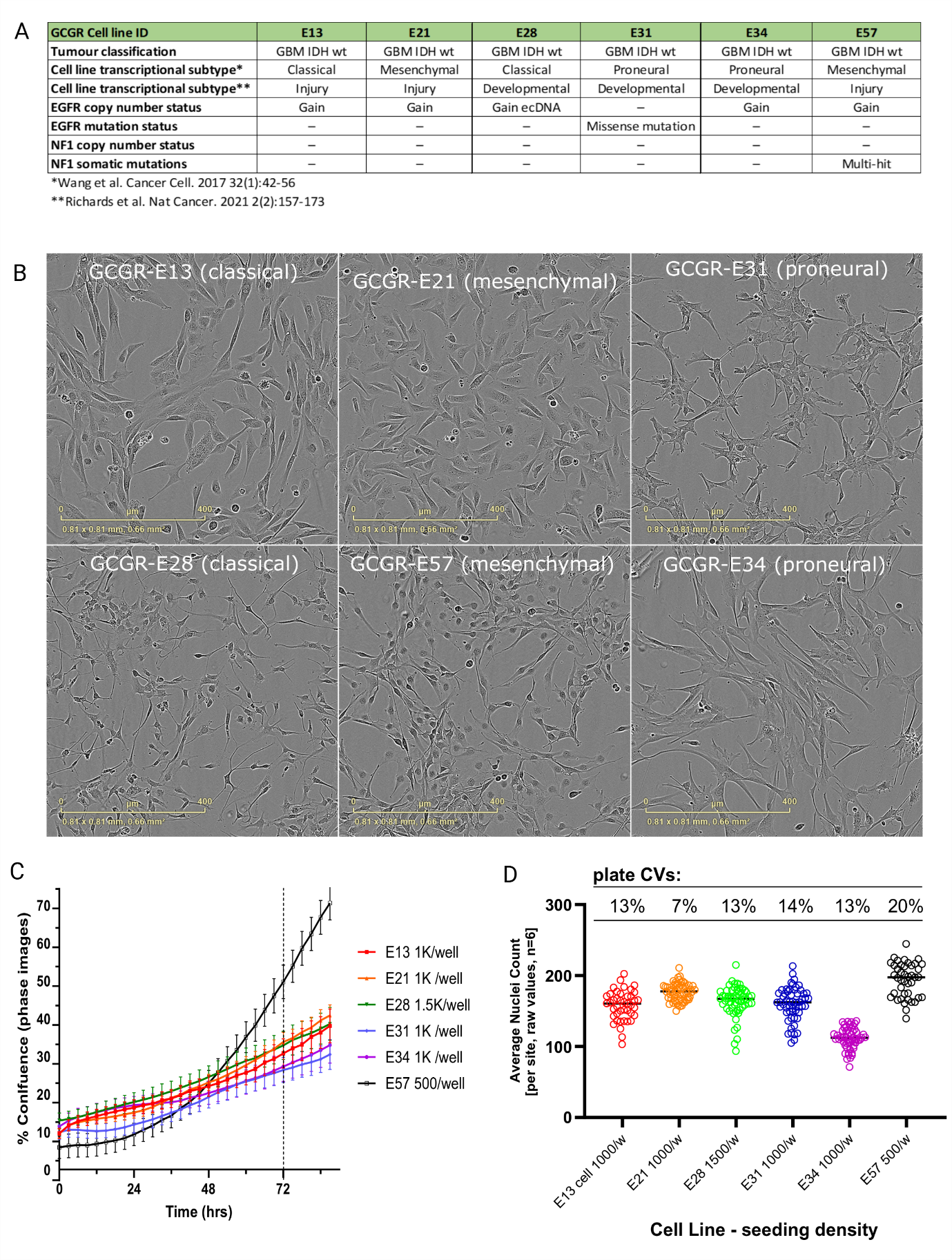

**Supplementary Figure 2. Basal characterisation of GSC cell models by reverse phase protein and cytokine array** A. Top ranked Z scores of normalised protein levels (268 antibodies inc. controls), with comparison to 2D and 3D conditions. Hierarchical clustering by one minus Pearson correlation and complete linkage. B. Network analysis of basal expression across all cell lines (2D+3D). C. Top ranked Z scores of basal cytokine expression in GSC models. D Network analysis of high basal expression of cytokines in GCGR cells.

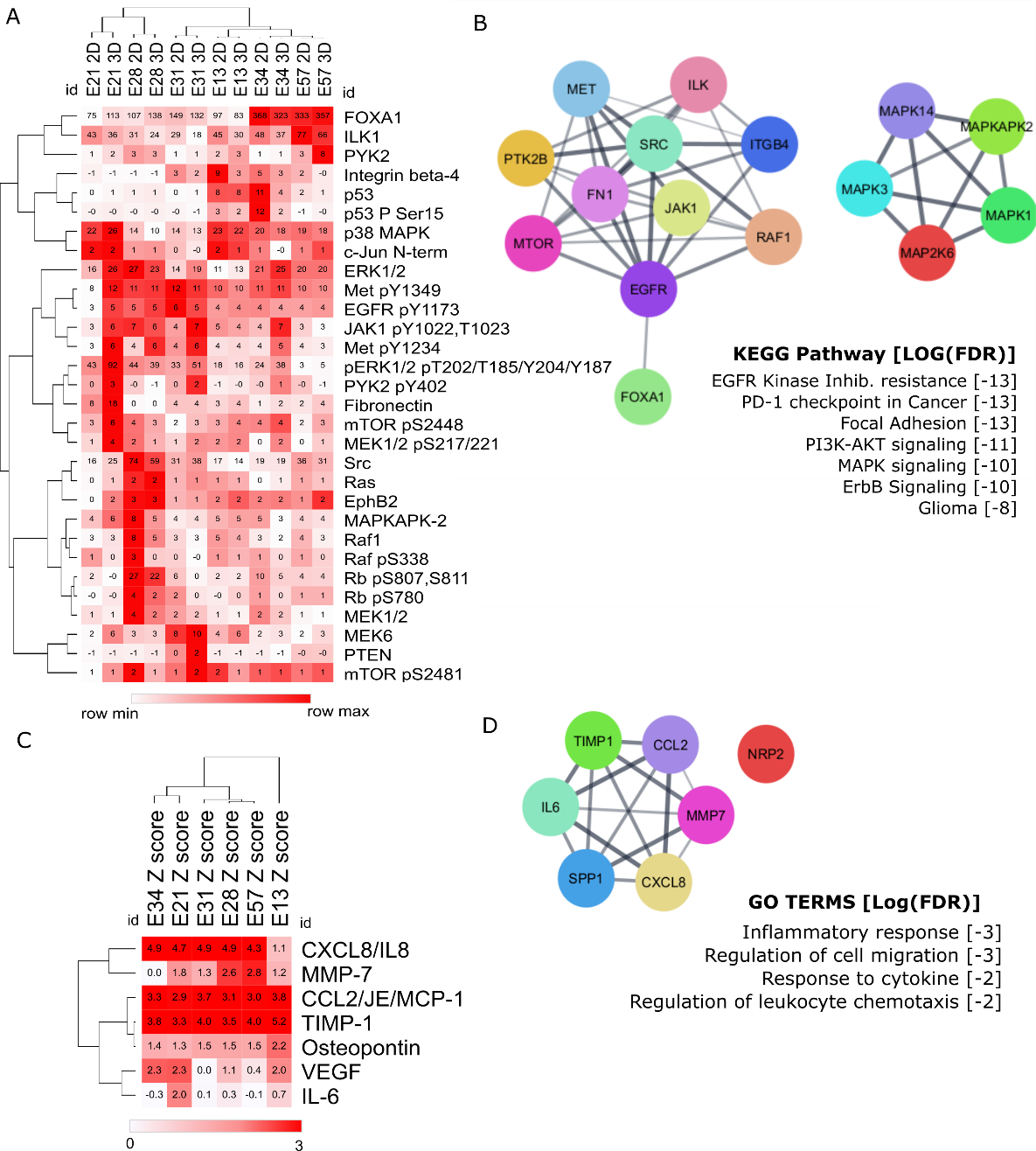

**Supplementary Figure 3**: Dose response optimisation of cell death on GCGR cells (positive controls, cell death) with IC50 values (nM) and Z prime calculations for 1µM doses.

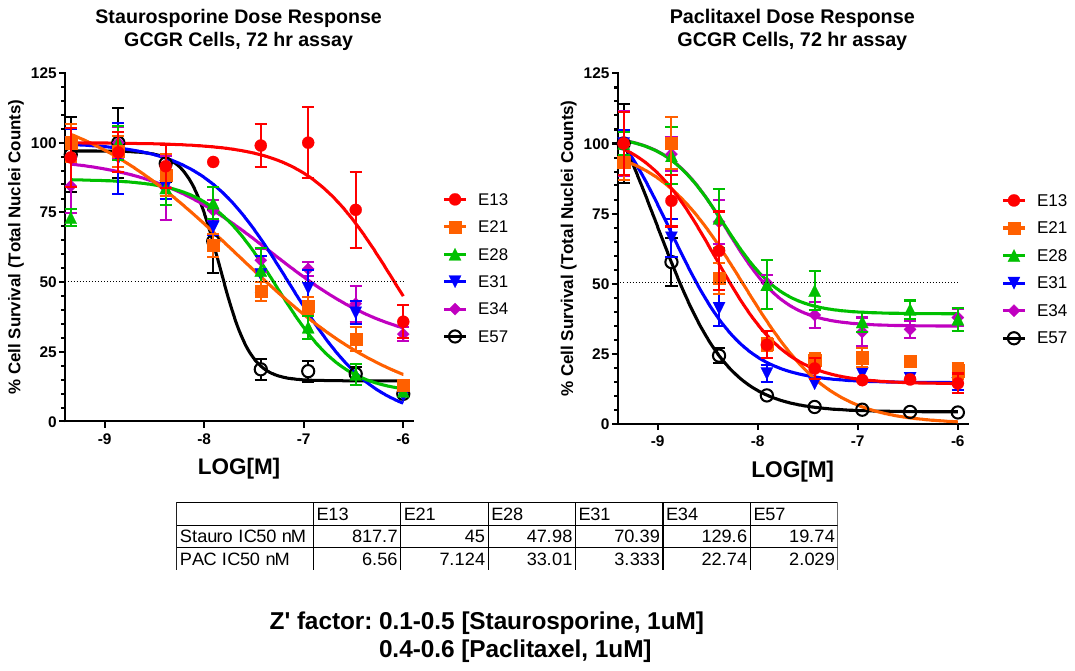

**Supplementary Figure 4: Pilot screening data**: TargetMol anti-cancer set L2110, 330 compounds (KCGS data excluded): A. Principal component analysis showing strong phenotypes in red (arbitrary thresholds) and B. Phenotypic distance vs Cell survival (normalised z score). Strongest hits in red, strong phenotypes in purple.

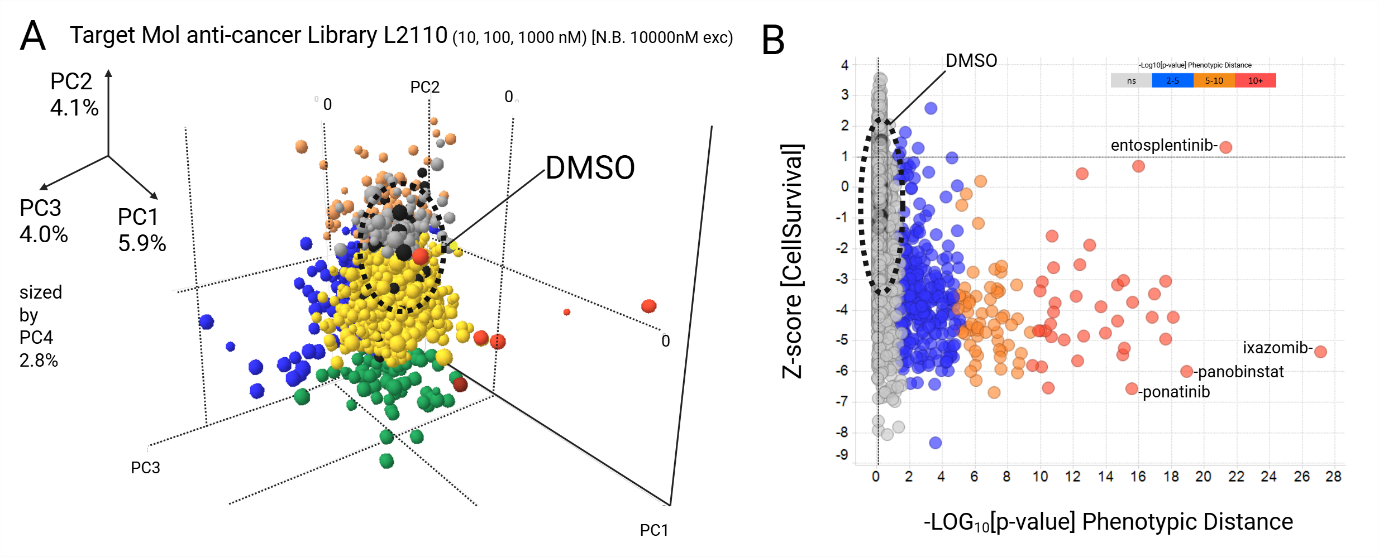

**Supplementary Figure 5.** Multiparametric phenotypic data analysis of compounds that affect cell viability and phenotypic distance far from DMSO controls[54]. (A, B) Cluster structure of the screened compounds with Z-Score [cell survival] < -3  and -LOG_10_[p-value] > 2 using the principal components derived from morphological features (Figure 2). Plot (A) shows the *k*-means clustering score and plot (B) the silhouette coefficient[55] averaged across compounds for an increasing number of clusters (*k*). Error bars denote one standard deviation over 125 repeats with random initial seeds. The lack of a clear “elbow” in the *k*-means score and low silhouette coefficients suggest poor clustering. (C) Distribution of -LOG_10_[p-value] Phenotypic Distance per chemical library and GCGR cell line screened, for the compounds with the same thresholds as in panels (A) and (B). The colour gradient quantifies -LOG_10_ [p-value] > 2. The heterogeneity of the cell lines is reflected in the diversity of the distributions observed, particularly in the mesenchymal cell lines (GCGR E21 and GCGR-E57).

 
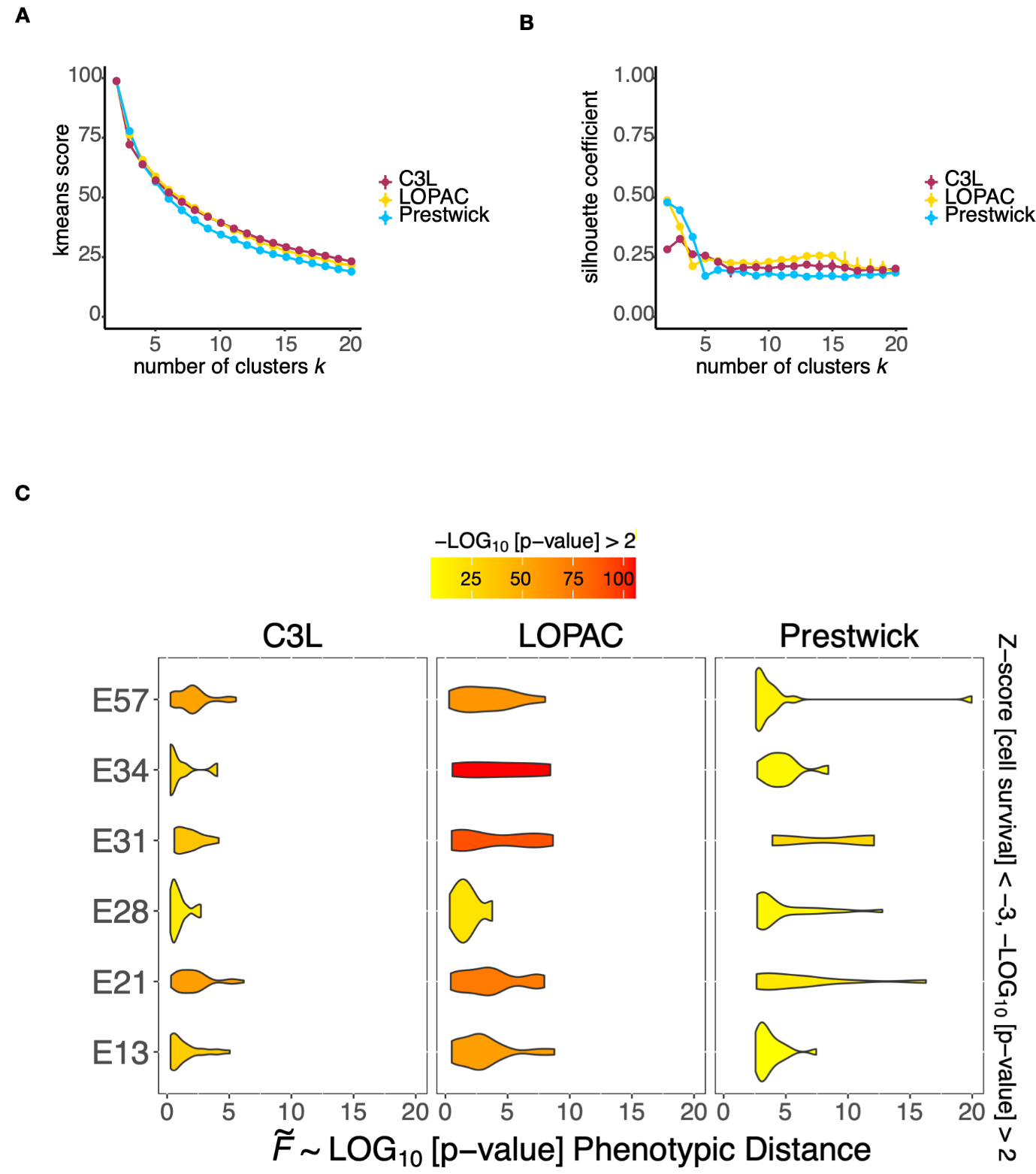

**Supplementary Figure 6.** STRING-Cytoscape network & enrichment analysis, of the annotated targets of the validated hits, demonstrates broad target-pathway coverage, including PI3K-AKT, MAPK, EGFR & adhesion signalling. B-F. Specific signalling pathways linked to annotated compound targets (highlighted in yellow) with False Discovery Rate [Log[FDR]] ranked GO terms.

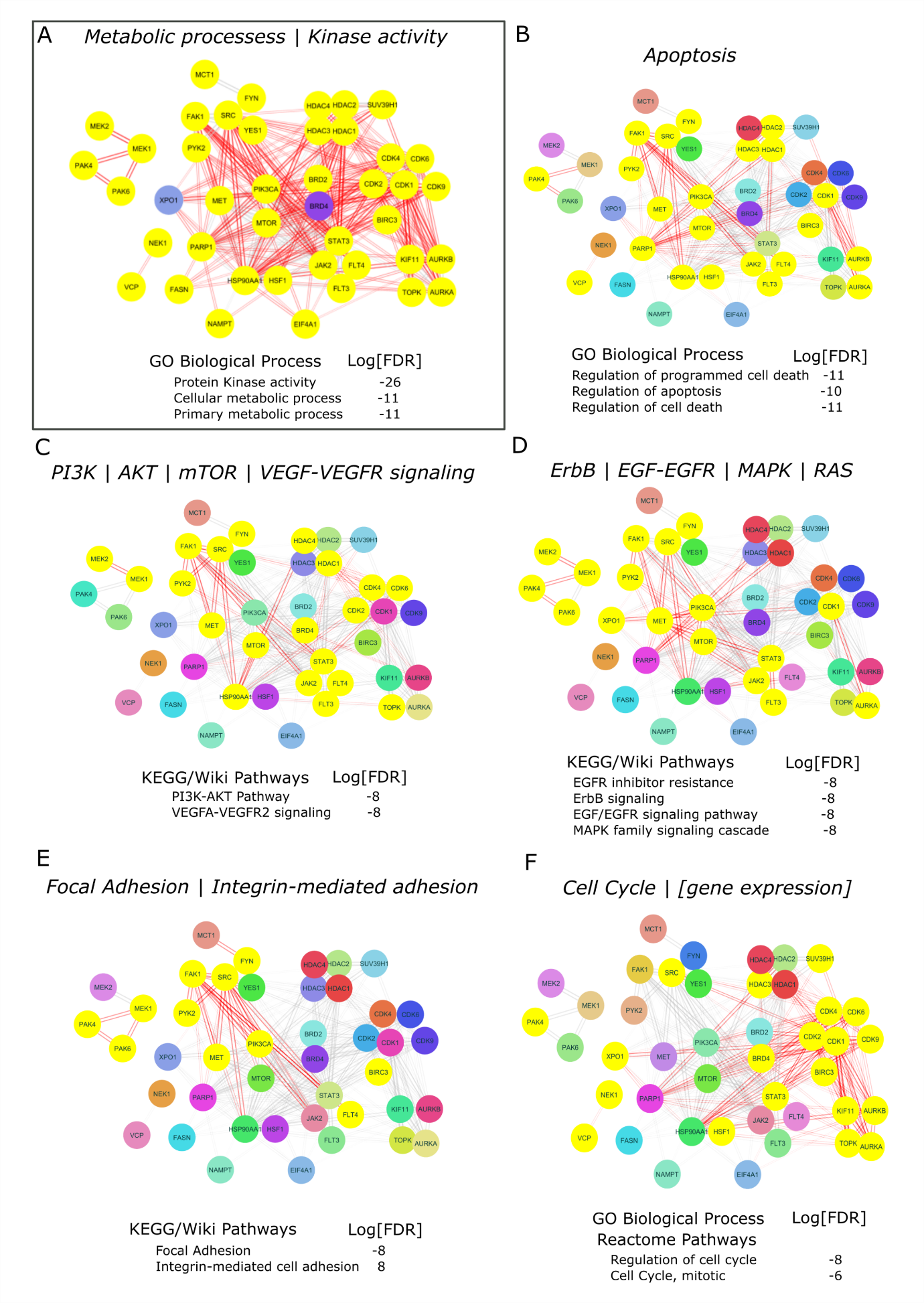

**Supplementary Figure 7**: Cell Cycle effects of HDAC and CDK9 inhibitors by dose response (%G0/G1 and (%G2/M) plus normalised nuclei counts). Calculated from nuclei stain/images & DNA content analysis.

A. Fimepinostat/CUDC-907 dose response (300nM to 0.6nM) across 6 GCGR cell lines. Percentage cell cycle (left y-axis) with %G0/G1 (red) and %G2/M (blue) with normalised nuclei counts as % cell survival (right y-axis, black). *[cell cycle data excluded where high doses result in complete cell death]*

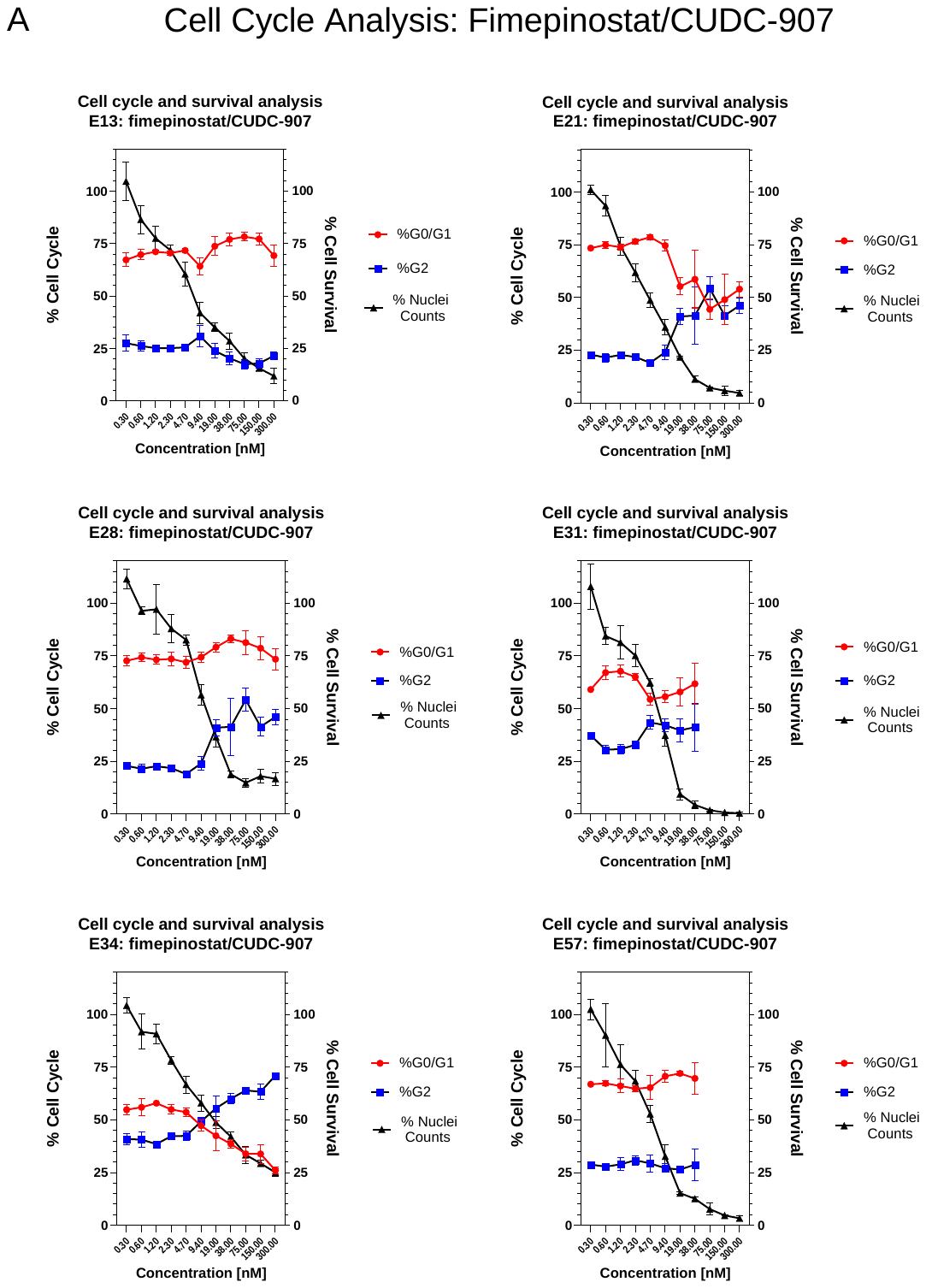

**Supplementary Figure 7 cont’d** : Cell Cycle effects of HDAC and CDK9 inhibitors by dose response (%G0/G1 and (%G2/M) plus normalised nuclei counts). Calculated from nuclei stain/images & DNA content analysis.

B. Romidepsin dose response (300nM to 0.6nM) across 6 GCGR cell lines. Percentage cell cycle (left y-axis) with %G0/G1 (red) and %G2/M (blue) with normalised nuclei counts as % cell survival (right y-axis, black). *[cell cycle data excluded where high doses result in complete cell death]*

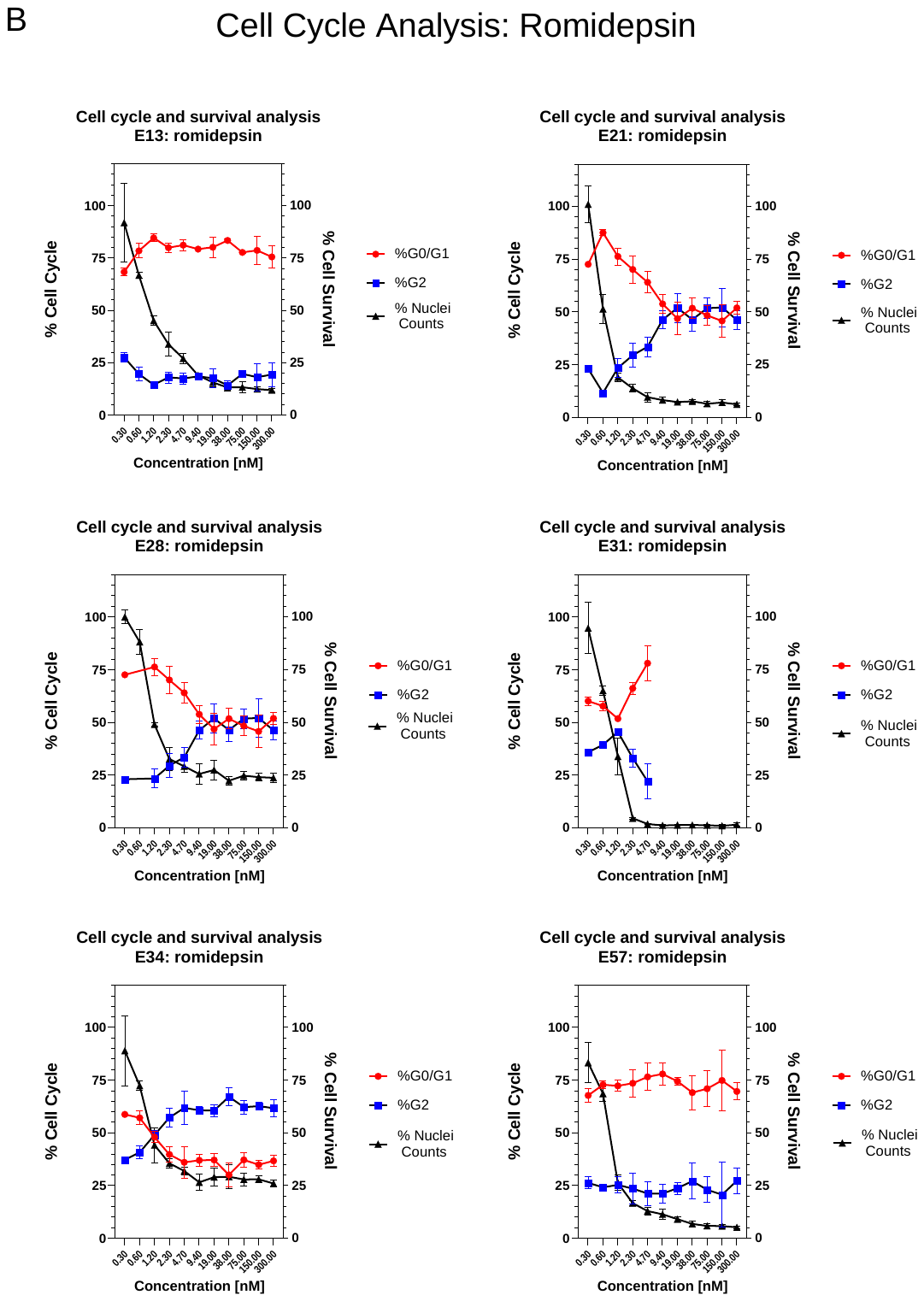

**Supplementary Figure 7 cont’d** : Cell Cycle effects of HDAC and CDK9 inhibitors by dose response (%G0/G1 and (%G2/M) plus normalised nuclei counts). Calculated from nuclei stain/images & DNA content analysis.

C. Panobinostat dose response (300nM to 0.6nM) across 6 GCGR cell lines. Percentage cell cycle (left y-axis) with %G0/G1 (red) and %G2/M (blue) with normalised nuclei counts as % cell survival (right y-axis, black). *[cell cycle data excluded where high doses result in complete cell death]*

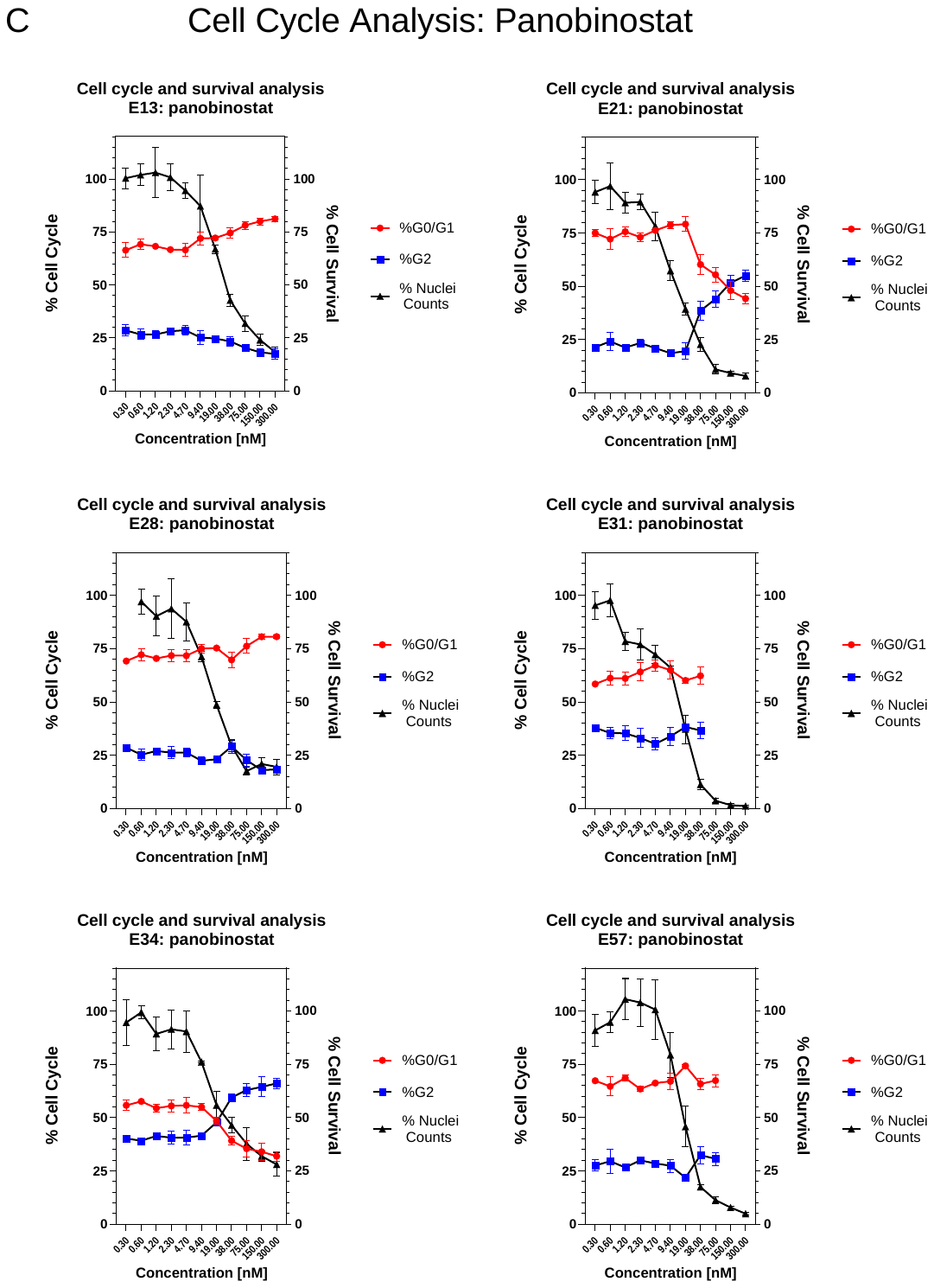

**Supplementary Figure 7 cont’d** : Cell Cycle effects of HDAC and CDK9 inhibitors by dose response (%G0/G1 and (%G2/M) plus normalised nuclei counts). Calculated from nuclei stain/images & DNA content analysis.

D. Dinaciclib dose response (300nM to 0.6nM) across 6 GCGR cell lines. Percentage cell cycle (left y-axis) with %G0/G1 (red) and %G2/M (blue) with normalised nuclei counts as % cell survival (right y-axis, black).

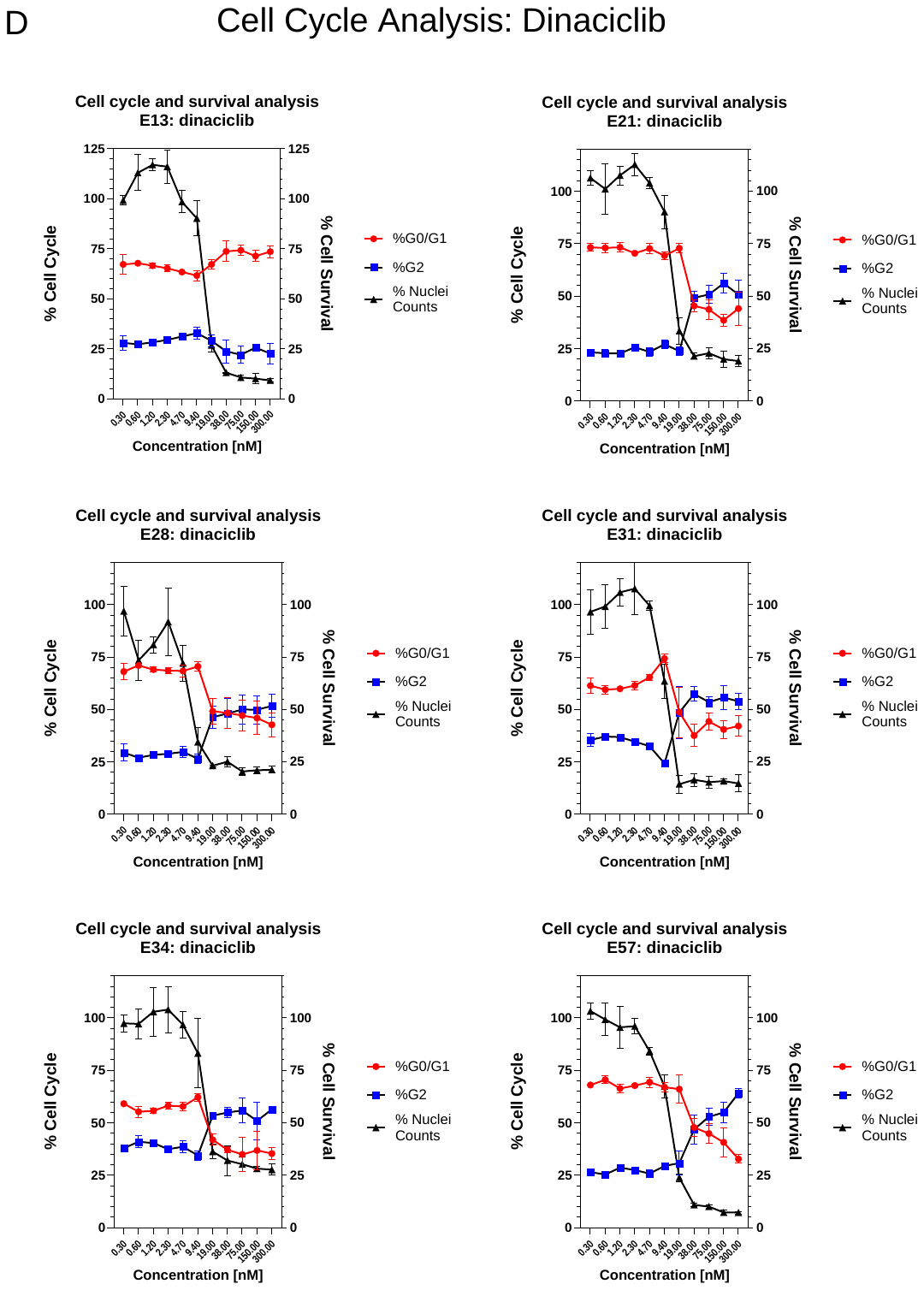

**Supplementary Figure 7 cont’d** : Cell Cycle effects of HDAC and CDK9 inhibitors by dose response (%G0/G1 and (%G2/M) plus normalised nuclei counts). Calculated from nuclei stain/images & DNA content analysis.

E. NVP-2 dose response (300nM to 0.6nM) across 6 GCGR cell lines. Percentage cell cycle (left y-axis) with %G0/G1 (red) and %G2/M (blue) with normalised nuclei counts as % cell survival (right y-axis, black).

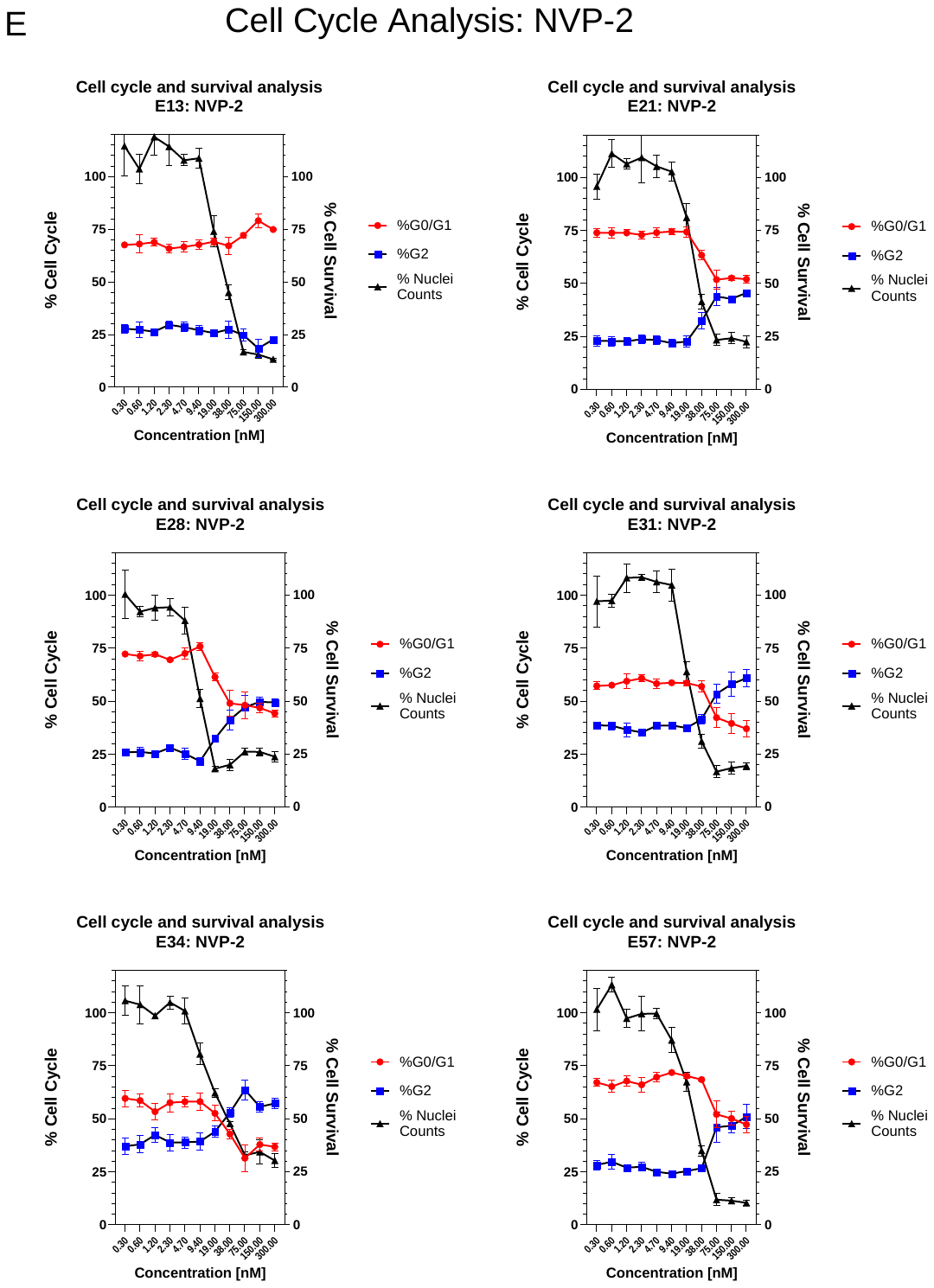

**Supplementary Figure 7 cont’d** : Cell Cycle effects of HDAC and CDK9 inhibitors by dose response (%G0/G1 and (%G2/M) plus normalised nuclei counts). Calculated from nuclei stain/images & DNA content analysis.

F. AZD4573 dose response (300nM to 0.6nM) across 6 GCGR cell lines. Percentage cell cycle (left y-axis) with %G0/G1 (red) and %G2/M (blue) with normalised nuclei counts as % cell survival (right y-axis, black).

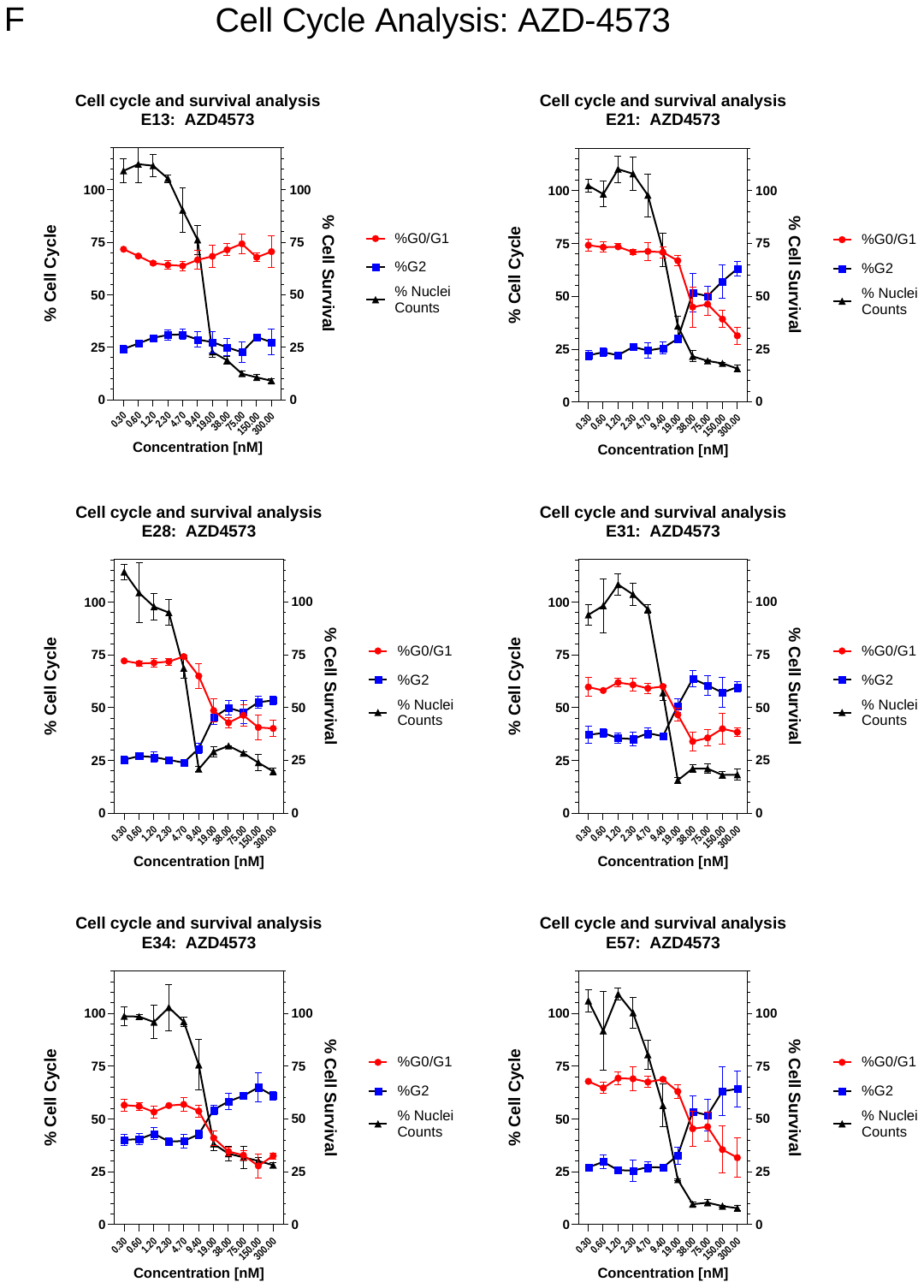

**Supplementary Figure 8**: Nanostring™ nCounter® transcriptomics and differential analysis of CDK9 inhibitors versus DMSO (0.1% (w/w)) on GCGR-E13 cells. A. Network enrichment analysis (string-db.org) of AZD4573, filtered by minimum 3-fold change and FDR <0.05 (277 genes). B. Venn diagram comparison of differentially expressed genes across AZD4573, dinaciclib & NVP-2. C. Differentially expressed genes unique to AZD4573 indicated. D. Summary of Biological processes (Gene Ontology) enriched in AZD4573 treated, GCGR-E13 cells (string-db.org). Coloured by FDR and sized by gene count.

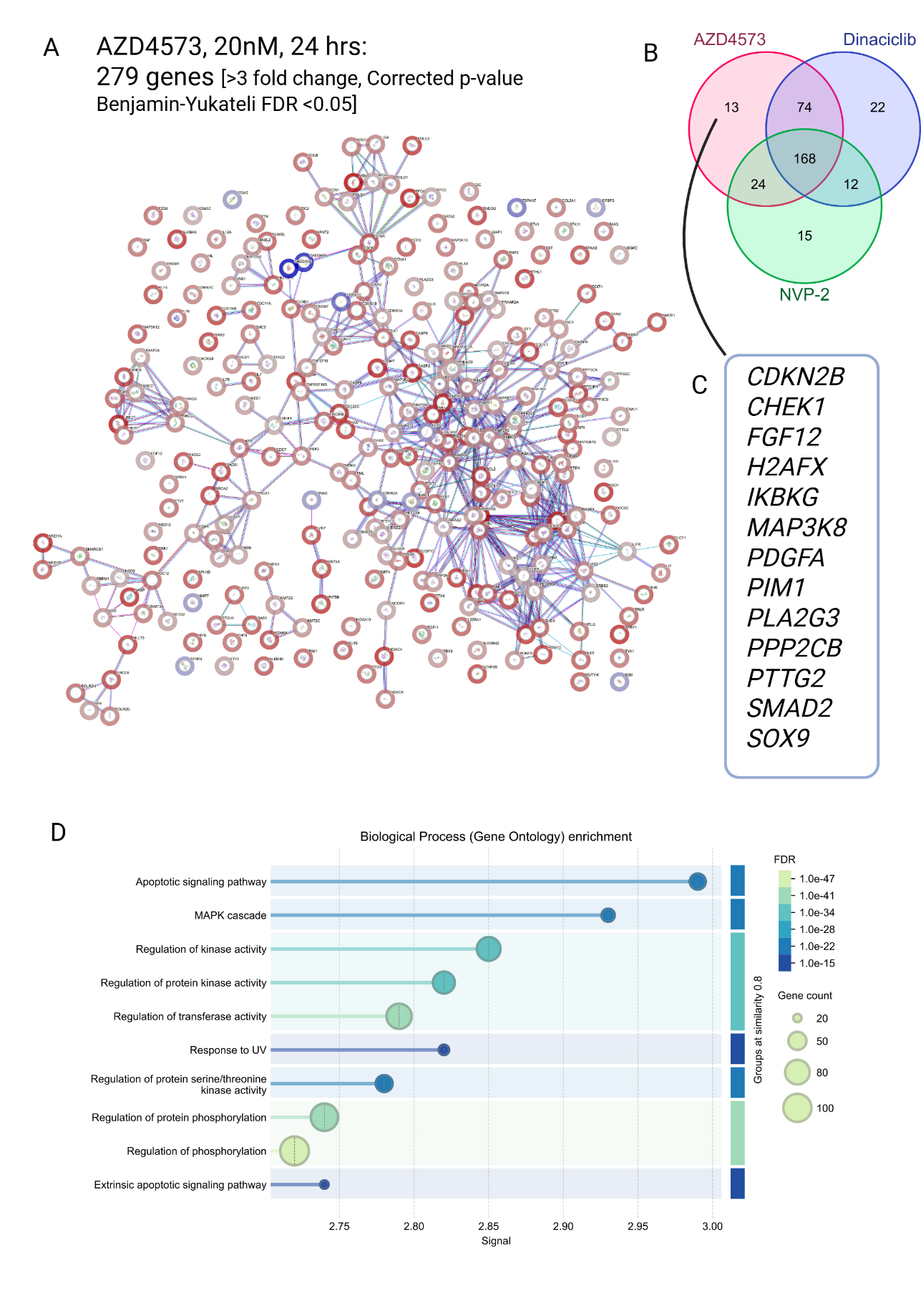

**Supplementary Figure 8 *continued***: Nanostring™ nCounter® transcriptomics and differential analysis of CDK9 inhibitors versus DMSO (0.1% (w/w)) on GCGR-E13 cells. E. Network enrichment analysis (string-db.org) of NVP-2, filtered by minimum 3-fold change and FDR <0.05 (277 genes). F. Venn diagram comparison of differentially expressed genes across AZD4573, dinaciclib & NVP-2 with differentially expressed genes unique to NVP-2 and (NVP-2 + AZD4573) indicated. G. Summary of Biological processes (Gene Ontology) enriched in NVP-2 treated, GCGR-E13 cells (string-db.org). Coloured by FDR and sized by gene count.

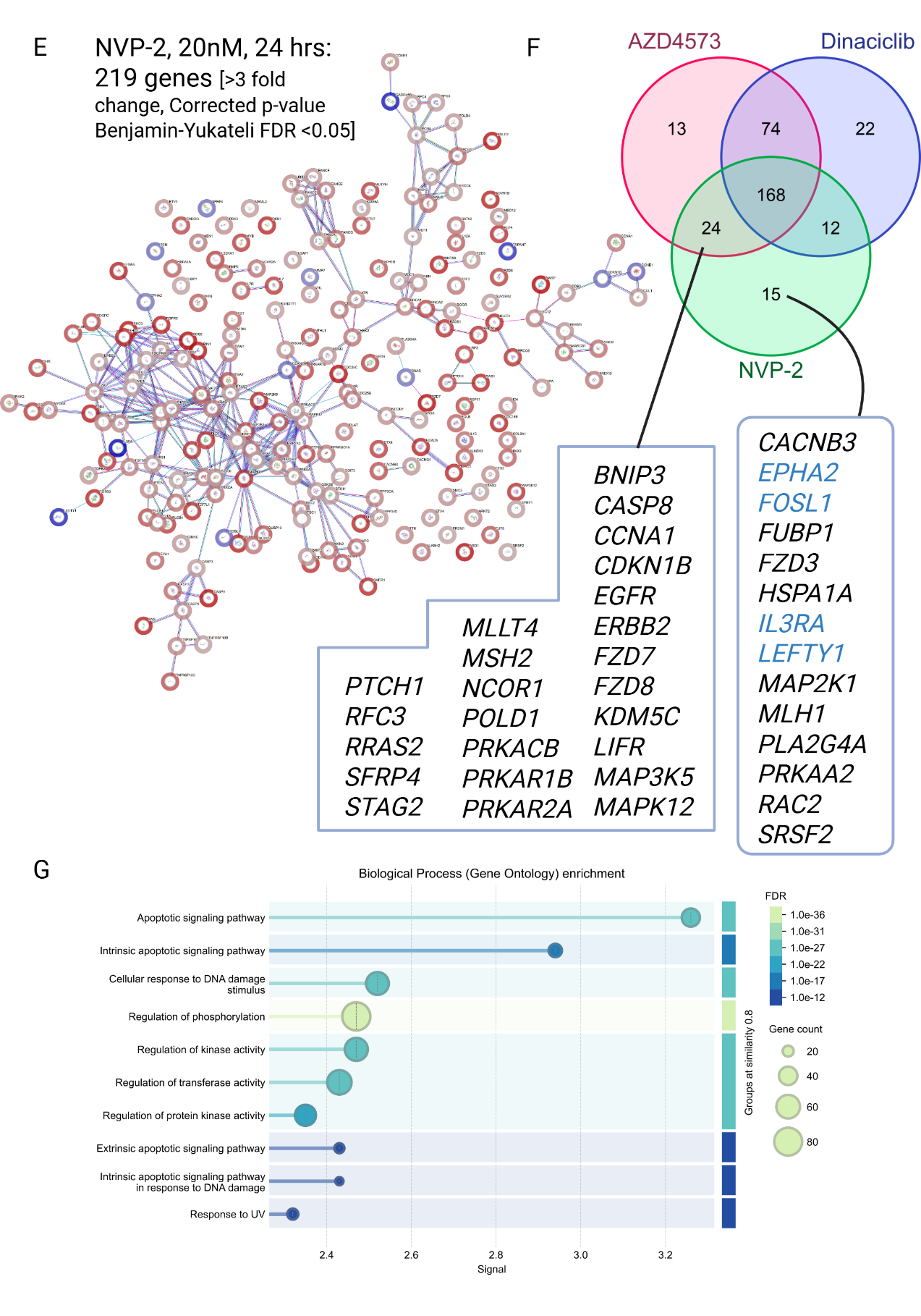

**Supplementary Figure 8 *continued***: Nanostring™ nCounter® transcriptomics and differential analysis of CDK9 inhibitors versus DMSO (0.1% (w/w)) on GCGR-E13 cells. H. Network enrichment analysis (string-db.org) of NVP-2, filtered by minimum 3-fold change and FDR <0.05 (277 genes) and Venn diagram comparison of differentially expressed genes across AZD4573, dinaciclib & NVP-2 with differentially expressed genes unique to dinaciclib indicated. I. Summary of Biological processes (Gene Ontology) enriched in dinaciclib treated, GCGR-E13 cells (string-db.org). Coloured by FDR and sized by gene count.

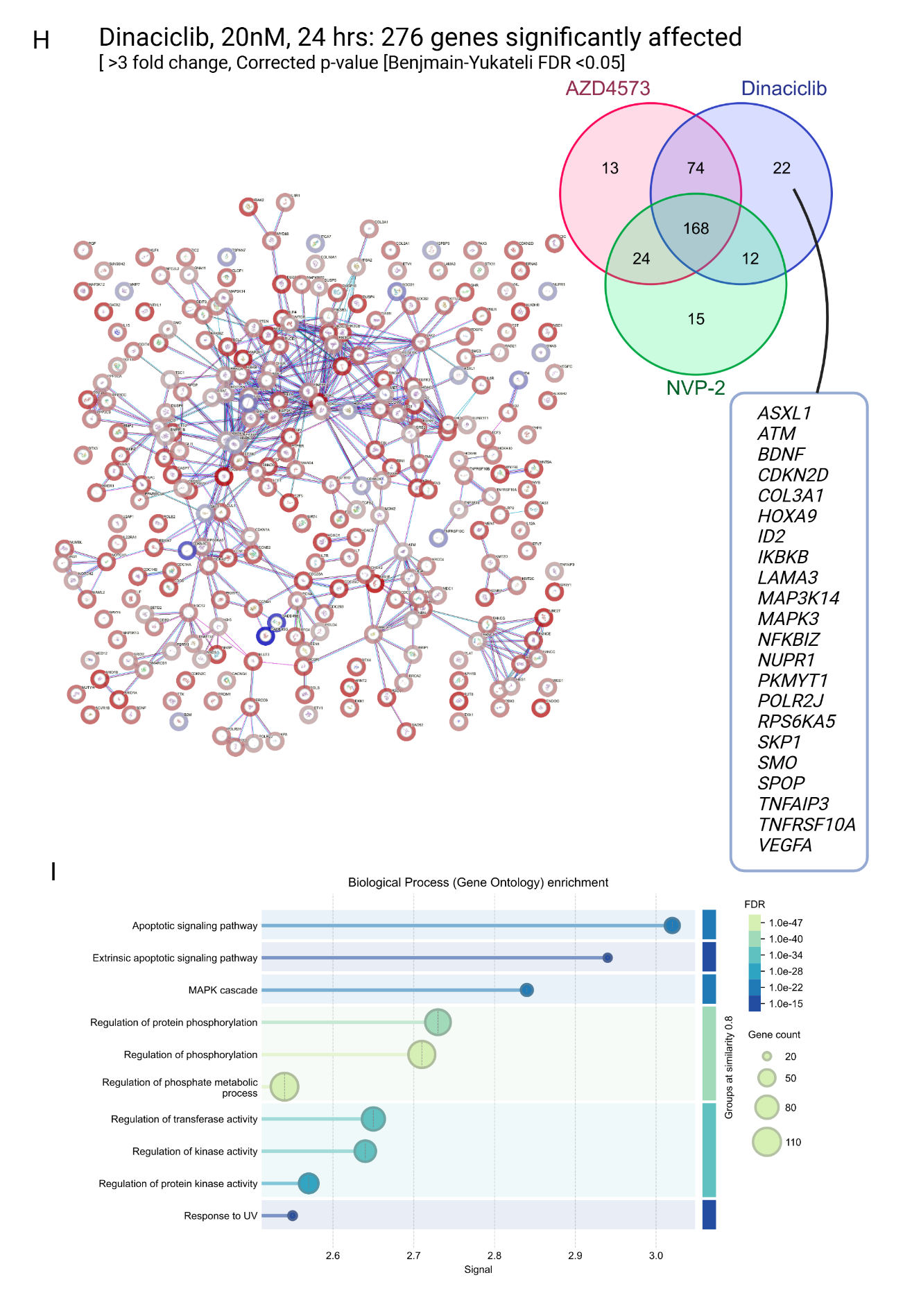

**Supplementary Figure 9. Drug combination experiments between Dinaciclib with romidepsin, idarubicin & ixazomib on GCGR-E13 cells**. A. % cell survival data of romidepsin across GCGR cells by dose response (reproduced from figure 4).B. DepMap correlation predicts effective combination with dinaciclib and romidepsin (R^2^ 0.657) across multiple cancer lineages (PRISM Repurposing Public 24Q2 dataset, depmap.org). C. Representative images of dinaciclib x romidepsin combination matrix (Hoechst and phalloidin stain (green)). Scale bar 100um. D. Combination matrix (% Inhibition max = red) of dinaciclib (25 – 0.5nM) and romidepsin (10 – 0.1nM) E. 3D Synergy plot of dinaciclib x romidepsin matrix (synergy score 75% quantile <10: no synergy observed).

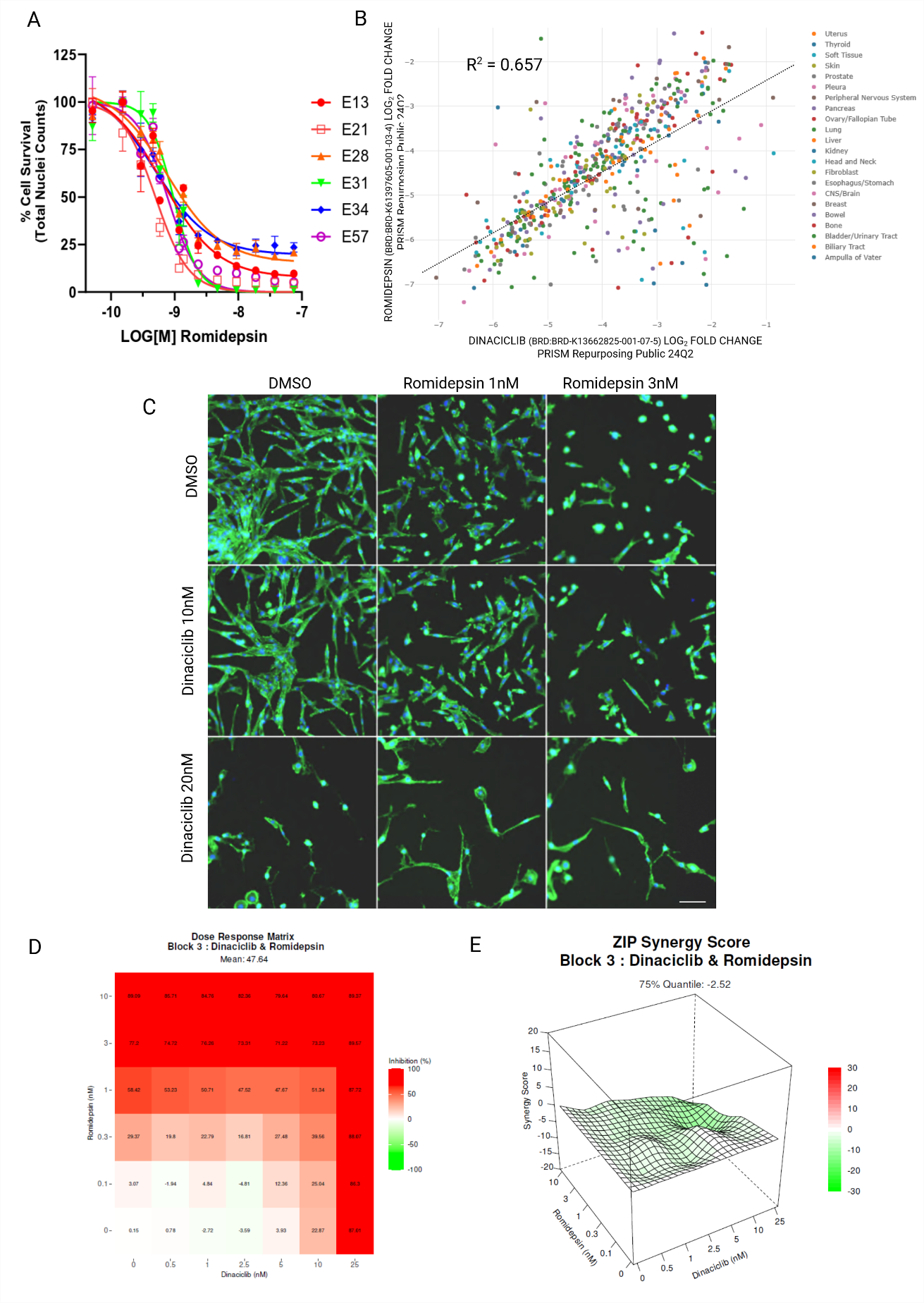

**Supplementary Figure 9 *continued*. Drug combination experiments between Dinaciclib with romidepsin, idarubicin & ixazomib on GCGR-E13 cells.** F. % cell survival data of idarubicin across GCGR cells by dose response. G. DepMap correlation predicts effective combination with dinaciclib and idarubicin (R^2^ 0.672) across multiple cancer lineages (PRISM Repurposing Public 24Q2 dataset, depmap.org). H. Representative images of dinaciclib x idarubicin combination matrix (Hoechst and phalloidin stain (green)). Scale bar 100um. I. Combination matrix (% Inhibition max = red) of dinaciclib (20 – 0.5nM) and idarubicin (80 – 1nM) J. 3D Synergy plot of dinaciclib x idarubicin matrix (synergy score 75% quantile <10: no synergy observed)

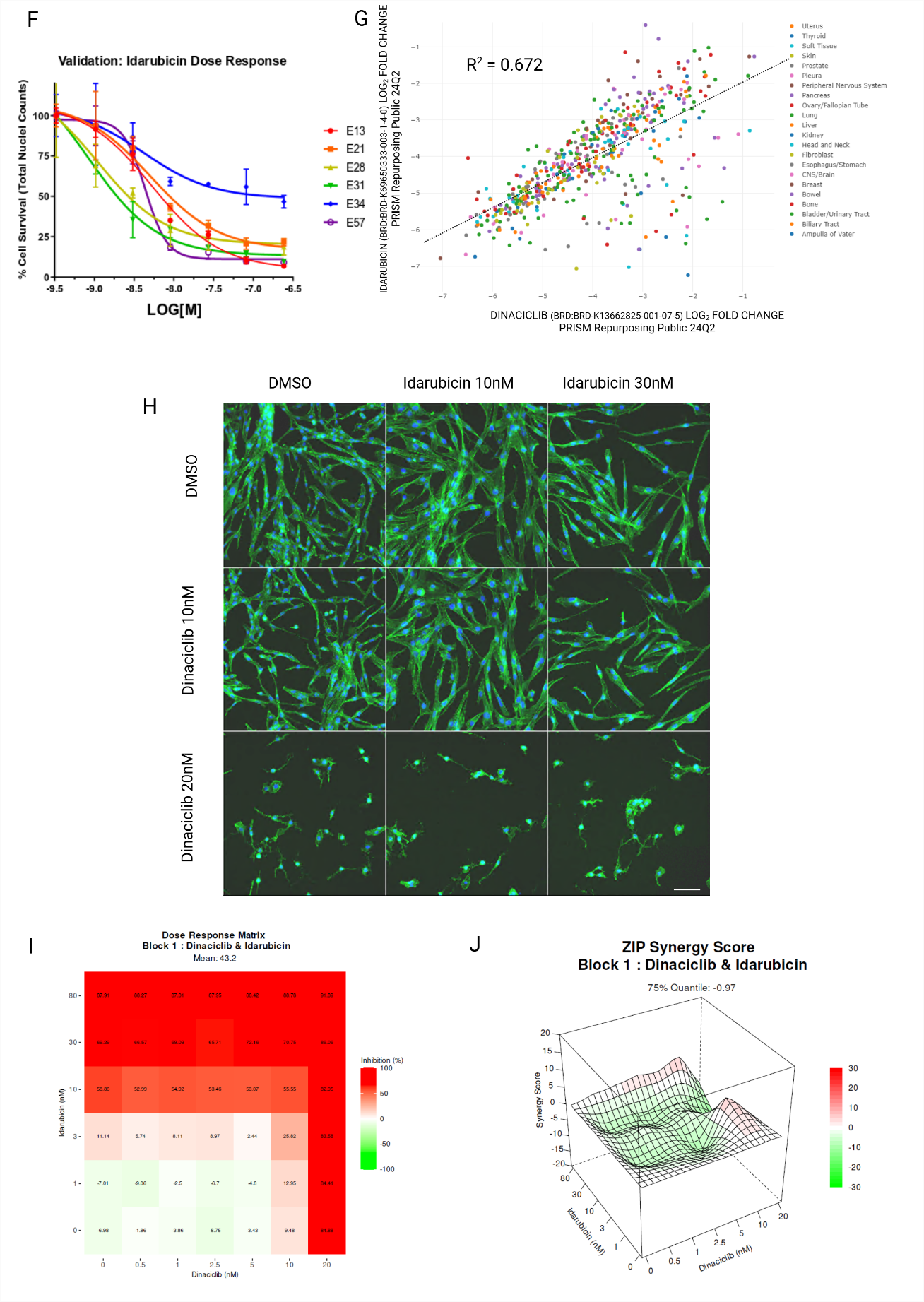

**Supplementary Figure 9 *continued.* Drug combination experiments between Dinaciclib with romidepsin, idarubicin & ixazomib on GCGR-E13 cells.** K. % cell survival data of ixazomib (citrate salt, MLN9708) across GCGR cells by dose response.. L. DepMap correlation predicts effective combination with dinaciclib and ixazomib (citrate salt, MLN9708) (R^2^ 0.622) across multiple cancer lineages (PRISM Repurposing Public 24Q2 dataset, depmap.org). M. Representative images of dinaciclib x ixazomib (citrate salt, MLN9708) combination matrix (Hoechst and phalloidin stain (green)). Scale bar 100um. N. Combination matrix (% Inhibition max = red) of dinaciclib (25 – 0.5nM) and ixazomib (citrate salt, MLN9708) (80 – 1nM) O. 3D Synergy plot of dinaciclib x ixazomib matrix (synergy score 75% quantile <10: no synergy observed)

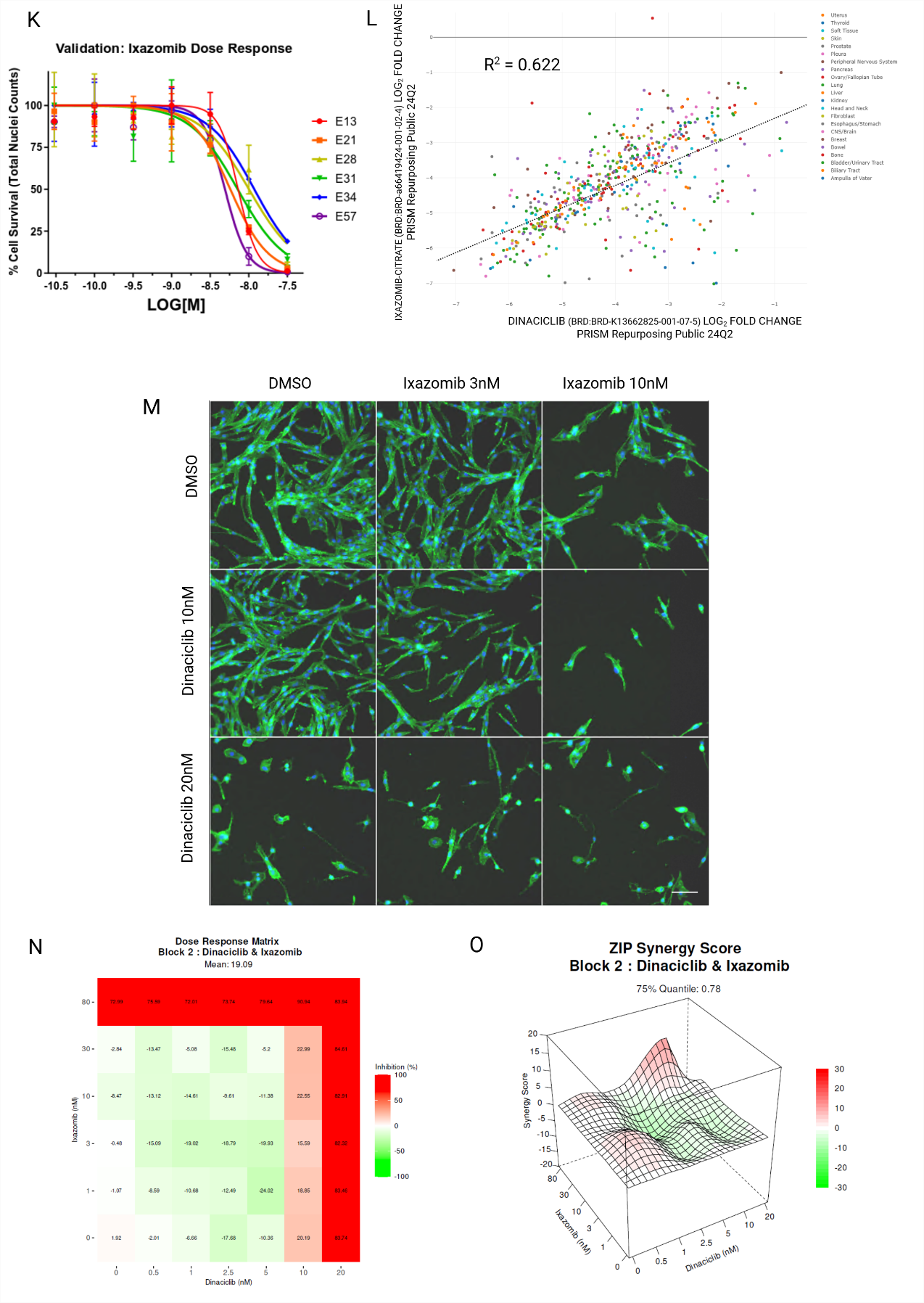

**Supplementary Figure 10. Comparison of commercial, potent PI3K inhibitors on E13 cells. A.** Live cell imaging of the effect of selective PI3K inhibitors versus dual PI3K/HDAC inhibitor, Fimepinostat (% confluence vs time (hrs) on GCGR-E13 cells. B. Summary table of published/quoted potencies (IC_50_ in nM) and selectivity across PI3K isoforms.

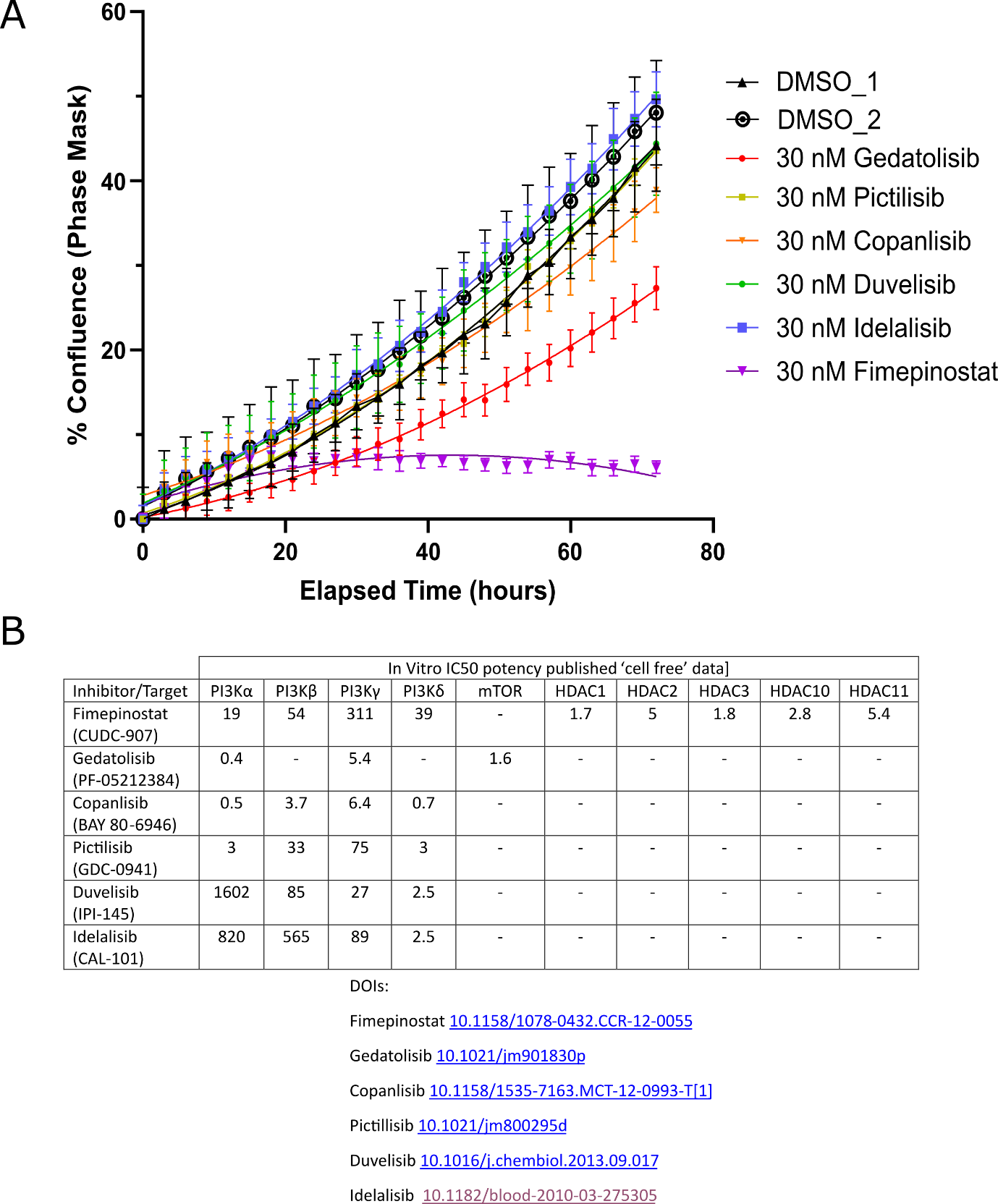

**Supplementary Table 1.** List of validated compounds showing biological activity up to 1µM in two or more glioma stem cell lines.

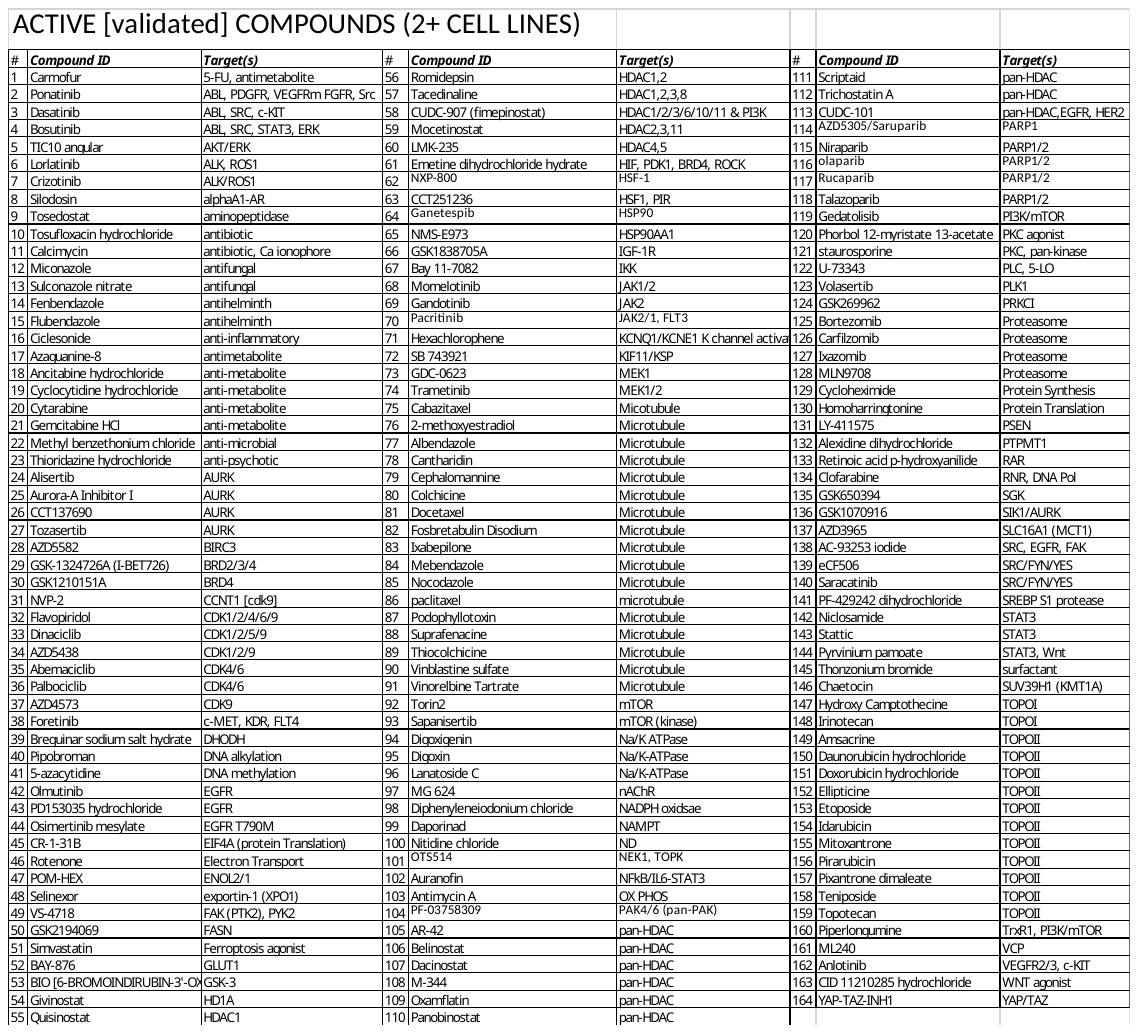
